# How short peptides can disassemble ultra-stable tau fibrils extracted from Alzheimer’s disease brain by a strain-relief mechanism

**DOI:** 10.1101/2024.03.25.586668

**Authors:** Ke Hou, Peng Ge, Michael R. Sawaya, Joshua L. Dolinsky, Yuan Yang, Yi Xiao Jiang, Liisa Lutter, David R. Boyer, Xinyi Cheng, Justin Pi, Jeffrey Zhang, Jiahui Lu, Shixin Yang, Zhiheng Yu, Juli Feigon, David S. Eisenberg

## Abstract

Reducing fibrous aggregates of protein tau is a possible strategy for halting progression of Alzheimer’s disease (AD). Previously we found that *in vitro* the D-peptide D-TLKIVWC disassembles tau fibrils from AD brains (AD-tau) into benign segments with no energy source present beyond ambient thermal agitation. This disassembly by a short peptide was unexpected, given that AD-tau is sufficiently stable to withstand disassembly in boiling SDS detergent. To consider D peptide-mediated disassembly as a potential therapeutic for AD, it is essential to understand the mechanism and energy source of the disassembly action. We find assembly of D-peptides into amyloid-like fibrils is essential for tau fibril disassembly. Cryo-EM and atomic force microscopy reveal that these D-peptide fibrils have a right-handed twist and embrace tau fibrils which have a left-handed twist. In binding to the AD-tau fibril, the oppositely twisted D-peptide fibril produces a strain, which is relieved by disassembly of both fibrils. This strain-relief mechanism appears to operate in other examples of amyloid fibril disassembly and provides a new direction for the development of first-in-class therapeutics for amyloid diseases.

## Introduction

Tau pathology caused by the abnormal aggregation of tau is more strongly correlated with cognitive symptoms and severity in Alzheimer’s disease (AD) than Aβ plaques^1–3^. Thus, disrupting tau aggregates emerges as a promising therapeutic strategy for AD^4,5^. To date, several types of disaggregators of tau fibrils have been reported and their disaggregation mechanisms are being sought. For example, the human Hsp70/DnaJ system can disaggregate tau fibrils *in vitro* in an ATP-dependent manner, but it generates small, seeding-competent species that accelerate the progression of disease^6^. Small molecules such as methylene blue^7^ and EGCG, by binding to tau fibrils, apparently disrupt intermolecular interactions^8^. Our recent structural studies revealed EGCG stacks in the clefts formed at the junction of the two protofilaments in AD tau fibrils^9^. However, small molecules including methylene blue/LMTX and EGCG have not been proven to be effective drugs, perhaps because of limited bioavailability, promiscuous protein binding and low bloodbrain barrier permeability^10^.

A peptide-based disaggregator can offer advantages over small molecules including higher specificity and higher binding affinity^11^. Especially, compared to L-peptides, D-enantiomeric peptides are known to be less immunogenic, less protease-sensitive *in vitro* and more resistant to degradation *in vivo*^12^. A recent phase I clinical trial of a D-peptide that disassembles Aβ oligomers has proved to be safe and well tolerated^13^. Previously, we reported that the D-peptide D-TLKIVWC can disassemble tau fibrils extracted from AD brains (AD-tau), neutralizing their seeding ability and rescuing behavioral deficits in a mouse model of Alzheimer’s disease^14^. However, the underlying disassembly mechanism remains unknown, preventing further development of this type of drug candidate for AD.

Here, we designed a series of peptides of sequence D-TLKIVWX varying only at the seventh residue, X. These D-peptides showed variable efficacy in disassembling AD-tau fibrils *in vitro*, with X = Ile as the best performer. From electron microscopy, we discovered that D-TLKIVWX peptides form amyloid-like fibrils themselves, and from atomic force microscopy we learned that these fibrils have a right-handed helical twist, in contrast to the left-handed helical twist of AD-tau. To learn the molecular interactions between the fibrils that participate in disassembly, we determined the cryo-EM structures of D-TLKIVWX protofilaments bound to tau fibrils of opposing twist. Combining our structural data, we propose a strain-relief mechanism for AD-tau fibril disassembly.

### D-TLKIVWX peptides disassemble AD-tau fibrils to non-seeding species

In previous work, we found that D-TLKIVWC can disassemble tau fibrils, whereas its six-residue analog D-TLKIVW cannot^14^. Here we wondered whether the disassembling ability arises from the capacity of the cysteine to form a disulfide bond with tau residue Cys-322 located in the structured core of tau fibrils. The formation of a disulfide bond could alter the conformation of tau in fibrils, resulting in fibril breakdown^15,16^. To evaluate this hypothesis, we measured disassembly activity as a function of disulfide bond forming potential which we adjusted by including either glutathione (GSH) or glutathione disulfide (GSSG). As shown in Extended Data Fig. 1a, D-TLKIVWC exhibited equivalent capacity to disassemble recombinant tau K18+ (residues Gln244-Glu380 of 4R tau) fibrils across all tested conditions, indicating the formation of a disulfide bond is irrelevant to the disassembly of tau fibrils.

To determine if the cysteine residue of D-TLKIVWC is essential in disassembling tau fibrils, we substituted the D-cysteine with various other D-amino acid residues, including anionic residues (aspartate (D) and glutamate (E)), cationic residues (arginine (R) and lysine (K)), polar residues (serine (S) and threonine (T)), hydrophobic residues (alanine (A), isoleucine (I), and valine (V)), as well as a β-sheet interrupter (proline (P))^17^. Their performance in disassembling AD-tau fibrils after 48 hours of incubation was evaluated by dot blot and electron microscopy (TEM). The dot blot experiment was conducted with the conformational antibody GT38^18^, which specifically recognizes AD-tau fibrils and does not probe monomeric tau or D-peptide controls (Extended Data Fig. 1b). The D-peptide variants (D-TLKIVWX, X = A, S, D, I, V, R, K, E, T) showed varying efficacy in disassembling AD-tau fibrils depending on the type of residue in the seventh position (Fig. 1a-b). Specifically, hydrophobic residues (X = I, V and A) most significantly reduced the level of AD-tau fibrils, with efficacy decreasing in the following order: hydrophobic > polar (X = S and T) > cationic (X = R and K) > anionic (X = D and E). Lastly, the β-sheet interrupter (X = P) and the deletion of the seventh residue both showed no reduction in the level of AD-tau fibrils. This trend was consistently supported by TEM characterization, where the number of AD-tau fibrils (labeled by red arrows in Fig. 1c and Extended Data Fig. 1c) was quantified after 48 hours of disassembly (Extended Data Fig. 1d). Notably, we observed new fibril species (blue arrows in Fig. 1c and Extended Data Fig. 1c). Furthermore, we observed specificity of D-TLKIVWX in disassembling AD-tau fibrils as it cannot disrupt other amyloid fibrils such as α-syn fibrils or wild type hnRNPA2 fibrils (Extended Data Fig. 2). In summary, most tested residue types X in D-TLKIVWX demonstrate specific disassembly action against AD-tau fibrils, with hydrophobic residues, especially Ile, being the best.

**Fig. 1:**
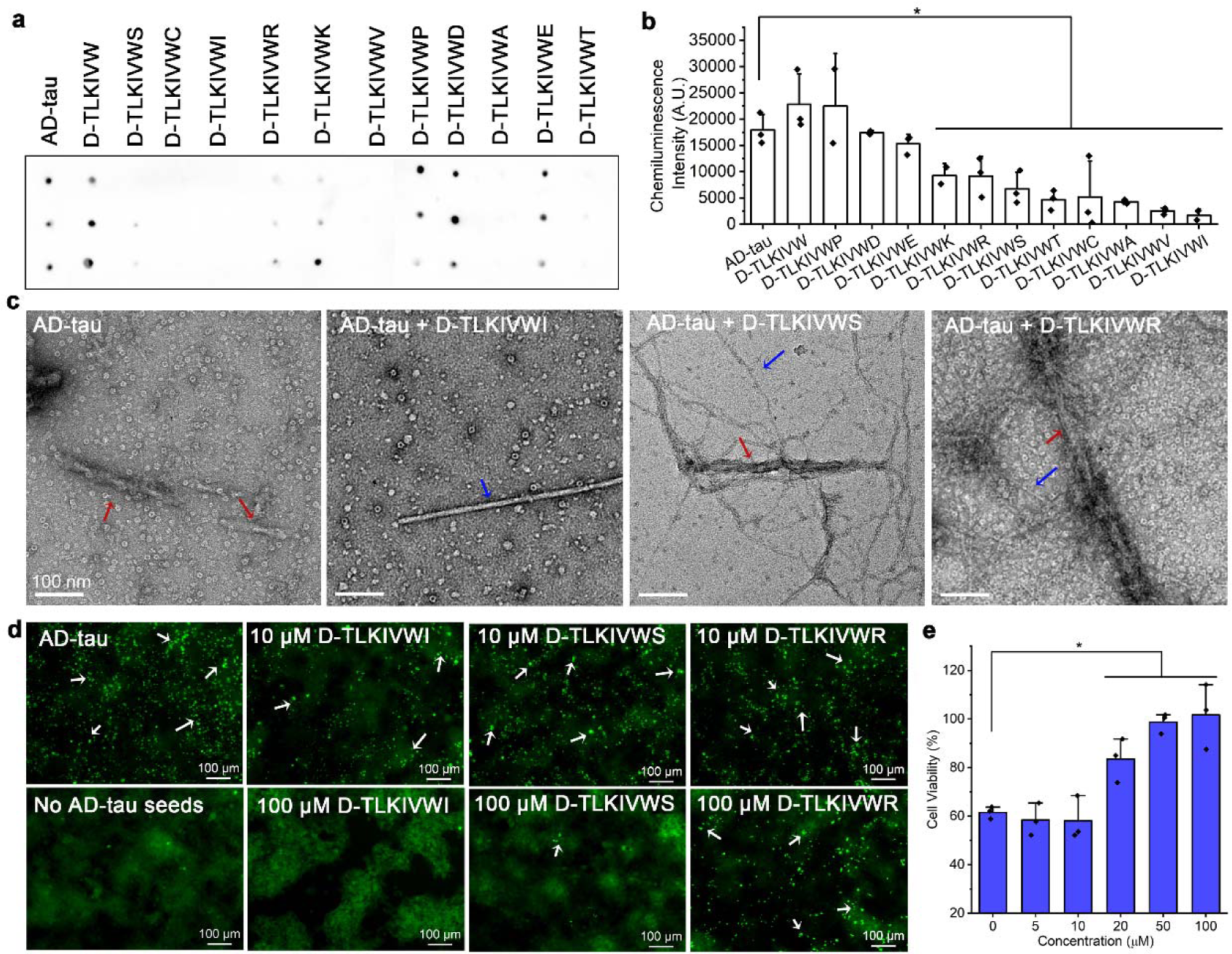
D-TLKIVWX peptides disassemble AD-tau fibrils to non-seeding products with varying efficiency. **a,** Dot blot staining of AD-tau fibrils before and after incubation with 500 μM D-TLKIVW, D-TLKIVWX (X = C, A, S, D, I, V, R, K, E, P, T) at 37 LJ for 48 h, probed by the GT38 antibody. Each sample was analyzed in triplicate. **b**, Quantification of the level of AD-tau fibrils in panel a by ImageJ. Statistical significance was analyzed by one-way ANOVA (\**p* < 0.05; the *p* vale is 0.026, 0.013, 0.002, 0.000, 0.001, 0.000, 0.000, 0.000 between AD-tau group and D-TKLIVWX (X = K, R, S, T, C, A, V, I) group, respectively). **c**, Representative TEM images of AD-tau fibrils after incubation with 500 μM D-TLKIVWX (X = I, S, R) at 37 LJ for 48 h. The AD-tau fibrils and newly formed fibrils are labeled by red and blue arrows, respectively. Scale bars, 100 nm. **d**, Representative fluorescent images of HEK293 cells expressing YFP-labeled tau K18 transfected with AD-tau fibril seeds after overnight disassembly with various concentrations of D-TLKIVWX (X = R, S, I). Fluorescent puncta are marked by white arrows. Scale bars, 100 μm. **e**, Cell viability of N2a cells treated with AD-tau fibrils in various concentrations of D-TLKIVWI measured by MTT assay. Results shown as Mean + SD of triplicate wells. Statistical significance was analyzed by one-way ANOVA (\**p* < 0.05; the *p* vale is 0.322, 0.021, 0.008, 0.009, 0.000 between 0 µM group and 5, 10, 20, 50 or 100 µM group, respectively).

In addition to uncovering the mechanism of disassembly, it is important to ascertain whether disassembly produces pathologic products that seed the growth of additional fibrils^19^. Therefore, we systematically investigated the seeding capacity of the products of AD-tau fibrils after overnight incubation with variants of D-TLKIVWX in a HEK293T cell line stably expressing yellow fluorescent protein (YFP)-fused tau^20^. As illustrated in Fig. 1d, AD-tau fibrils alone can seed the aggregation of endogenous fluorescent tau, leading to the formation of bright puncta (indicated as white arrows) observable under fluorescence microscopy. In contrast, the overnight disassembly products of AD-tau fibrils treated with D-TLKIVWX gradually lost their seeding ability in a dose-dependent manner (Fig. 1d and Extended Data Fig. 3a). Further, automated image analysis of visible puncta revealed a dependence of the seeding capacity on the seventh amino acid residue within D-TLKIVWX (Extended Data Fig. 3b-f). The observed dependence is consistent with the trends observed in our *in vitro* disassembly assays (Fig. 1b and Extended Data Fig. 1c,d). Additionally, because of disassembly activity, D-TLKIVWX (X = I and S) shows a dose-dependent effect in reducing AD-tau toxicity in mouse Neuro 2A (N2a) cells (Fig. 1e and Extended Data Fig. 3g). Our results demonstrate that the products of AD tau fibrils disassembled by D-TLKIVWX are not seeding-competent and are non-toxic.

### Amyloid-like fibril formation of D-TLKIVWX is essential for tau disassembly

To better understand the disassembly process of tau fibrils, we conducted a time-course dot blot and TEM experiment using D-TLKIVWI as a representative of the more efficient fibril disassemblers. Dot blot data (Extended Data Fig. 4a) confirmed that D-TLKIVWI gradually reduced the level of AD-tau fibrils as a function of time. The time-course TEM images in Fig. 2a and Extended Data Fig. 4b show AD-tau fibrils (red arrows) appearing to be covered by unknown species after one hour of incubation with D-TLKIVWI, and additional fibrillar structures (blue arrows) become increasingly evident at three-and six-hour timepoints. At 24 h, amorphous products (magenta arrows) appeared concomitant with the disappearance of AD-tau fibrils and reduction of the unidentified fibril species. The amorphous products appeared more dispersed in micrographs acquired at 48 h, and a western blot showed that the disassembly products of AD-tau fibrils consisted primarily of insoluble species; denatured pelleted material migrated as dimers and other multimers, not as monomeric tau (Extended Data Fig. 4c). Notably, in the TEM images of D-TLKIVWX (X = I, S, R, D, E, K, T, C, A and V)-treated AD-tau samples (Fig. 1c and Extended Data Fig. 1c), we also observed the emergence of new fibrils (labeled by blue arrows) accompanying with the disappearance of AD-tau fibrils. We identified these new species as D-TLKIVWX fibrils since D-TLKIVWX (X = I, S, R, D, E, K, T, C, A and V) exhibited aggregation activity in the same buffer (Extended Data Fig. 5a). In contrast, neither D-TLKIVW nor D-TLKIVWP exhibited aggregation nor disassembled AD-tau fibrils. These observations suggest that the ability of D-TLKIVWX to fibrilize aids the disassembly of AD-tau fibrils.

**Fig. 2:**
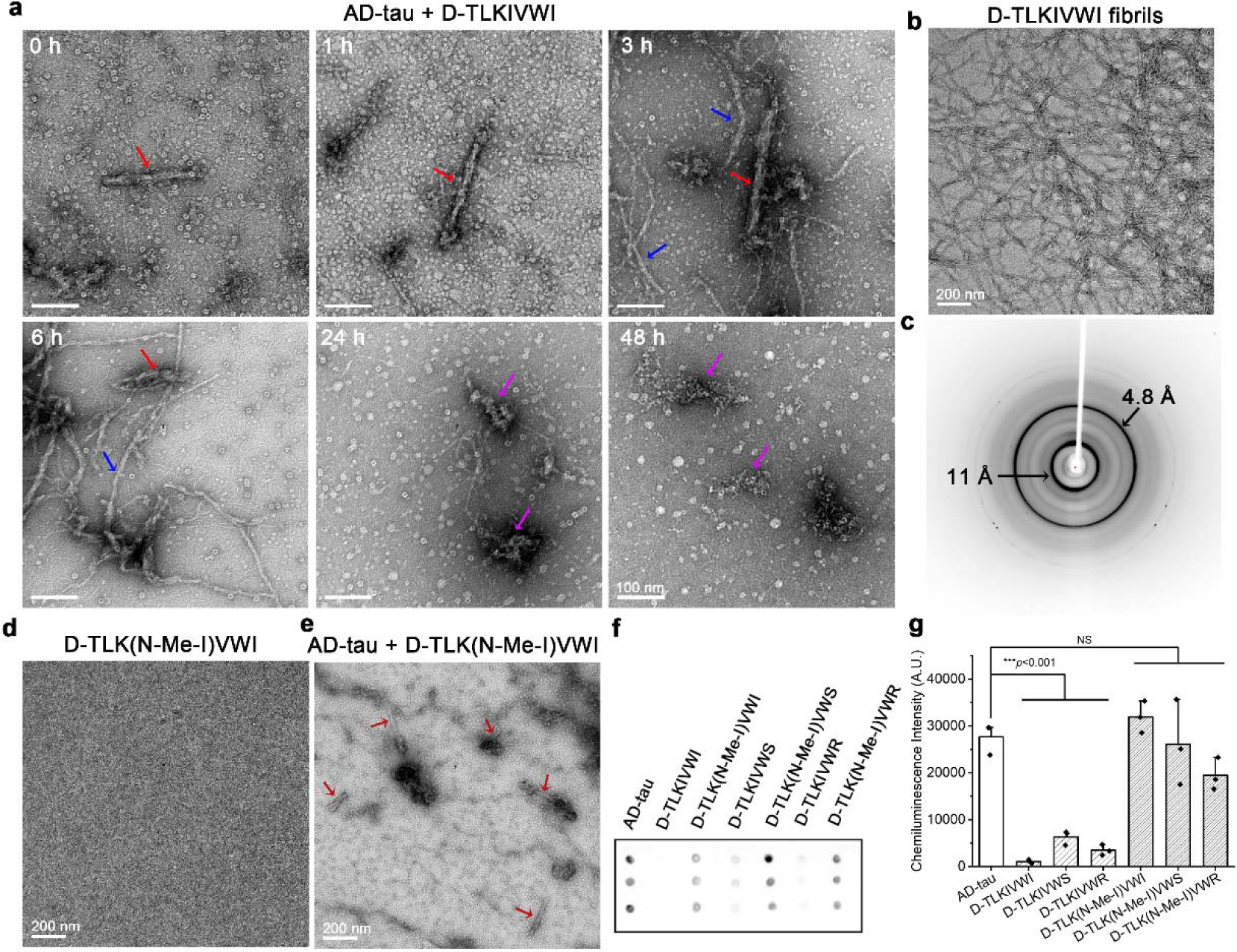
D-TLKIVWX disassembly of AD-tau fibrils involves formation of D-TLKIVWX amyloid-like fibrils. **a**, Negative stain transmission electron micrographs of AD-tau fibrils incubated with 500 μM D-TLKIVWI at 0, 1, 3, 6, 24 and 48 h timepoints. AD-tau fibrils, newly formed D-TLKIVWI fibrils, and amorphous products are labeled by red, blue and magenta arrows, respectively. **b**, Negative stain transmission electron micrograph of D-TKLIVWI amyloid-like fibrils formed by 10 mM D-TLKIVWI grown under quiescent conditions in deionized water at room temperature for three days. **c**, X-ray diffraction pattern of D-TLKIVWI fibrils showing spacings characteristic of amyloid-like fibrils. **d**, Negative stain transmission electron micrograph of 500 μM D-TLK(N-Me-I)VWI incubated in 20 mM Tris-HCl, pH 7.4, 100 mM NaCl at 37 LJ for 48 h. **e**, Representative TEM image of AD-tau fibrils after incubation with 500 μM D-TLK(N-Me-I)VWI at 37 LJ for 48 h. AD-tau fibrils are labeled by red arrows. **f**, Dot blot staining of AD-tau fibrils before and after incubation with 500 μM D-TLKIVWX or D-TLK(N-Me-I)VWX (X = I, S and R) at 37 LJ for 48 h, probed by the GT38 antibody. **g**, Quantification of the dot blot staining in panel f. Results shown as Mean + SD of triplicate dots. Statistical significance was analyzed by one-way ANOVA (\*\*\**p* < 0.001; NS, not significant; the *p* vale is 0.000, 0.000, 0.000, 0.869, 0.999, 0.263 between AD tau group and D-TLKIVWX (X= I, S, or R) or D-TLK(N-Me-I)VWX (X = I, S or R) group, respectively).

To test the hypothesis that D-TLKIVWX form amyloid-like fibrils that disassemble AD-tau, we designed a negative control experiment by eliminating the ability of D-TLKIVWX to fibrilize. As shown in Fig. 2b and Extended Data Fig. 5b,c, 10 mM D-TLKIVWX (X = I, S and R) form well-defined fibrils when they are left undisturbed for three days at room temperature. The corresponding X-ray diffraction analysis (Fig. 2c and insets in Extended Data Fig. 5b,c) confirm these D-TLKIVWX (X = I, S and R) fibrils exhibit the characteristic features of amyloid fibrils with a strong 4.7-4.8 Å reflection corresponding to the distance between hydrogen-bonded β-strands, and a more diffuse 8-12 Å ring arising from the inter-sheet spacing^21^. As such, the aggregation activity of D-TLKIVWX might be prevented by eliminating hydrogen bonds between neighboring β-strands through N-methylation of peptide backbones^22^. Indeed, when we N-methylated the D-isoleucine of D-TLKIVWX (named as D-TLK(N-Me-I)VWX (X = I, S and R)), the peptides were unbale to form fibrils, as shown in Fig. 2d and Extended Data Fig. 5d,e. TEM and dot blot experiments further showed that these non-self-aggregating peptides D-TLK(N-Me-I)VWX (X = I, S and R) were unable to disassemble AD-tau fibrils (Fig. 2e-g and Extended Data Fig. 5f,g). Taken together, these experiments demonstrate the critical role of amyloid-like characteristics of D-TLKIVWX in disassembling AD-tau fibrils.

### D-TLKIVWX peptides form right-handed fibrils with conserved motifs

We determined the structures of D-TLKIVWX amyloid-like fibrils, aiming to reveal features that facilitate disassembly of AD-tau fibrils. Generally, β-sheets formed by L-peptides adopt left-handed helical structures, while β-sheets formed by D-peptides adopt right-handed helical structures^23,24^. Here, atomic force microscopy (AFM) confirmed the twist is right-handed in all 18 polymorphs of D-TLKIVWI fibrils observed^25,26,27^ (Fig. 3a and Extended Data Fig. 6a). Using cryo-EM, we were able to determine the structures of the predominant polymorphs of D-TLKIVWX (X = I, S and R) fibrils (indicated by red squares in Extended Data Fig. 6b-d) at 3.6 Å, 3.5 Å, 3.7 Å resolution, respectively (Fig. 3b-d). Data collection and refinement statistics are summarized in Supplementary Table 1.

**Fig. 3:**
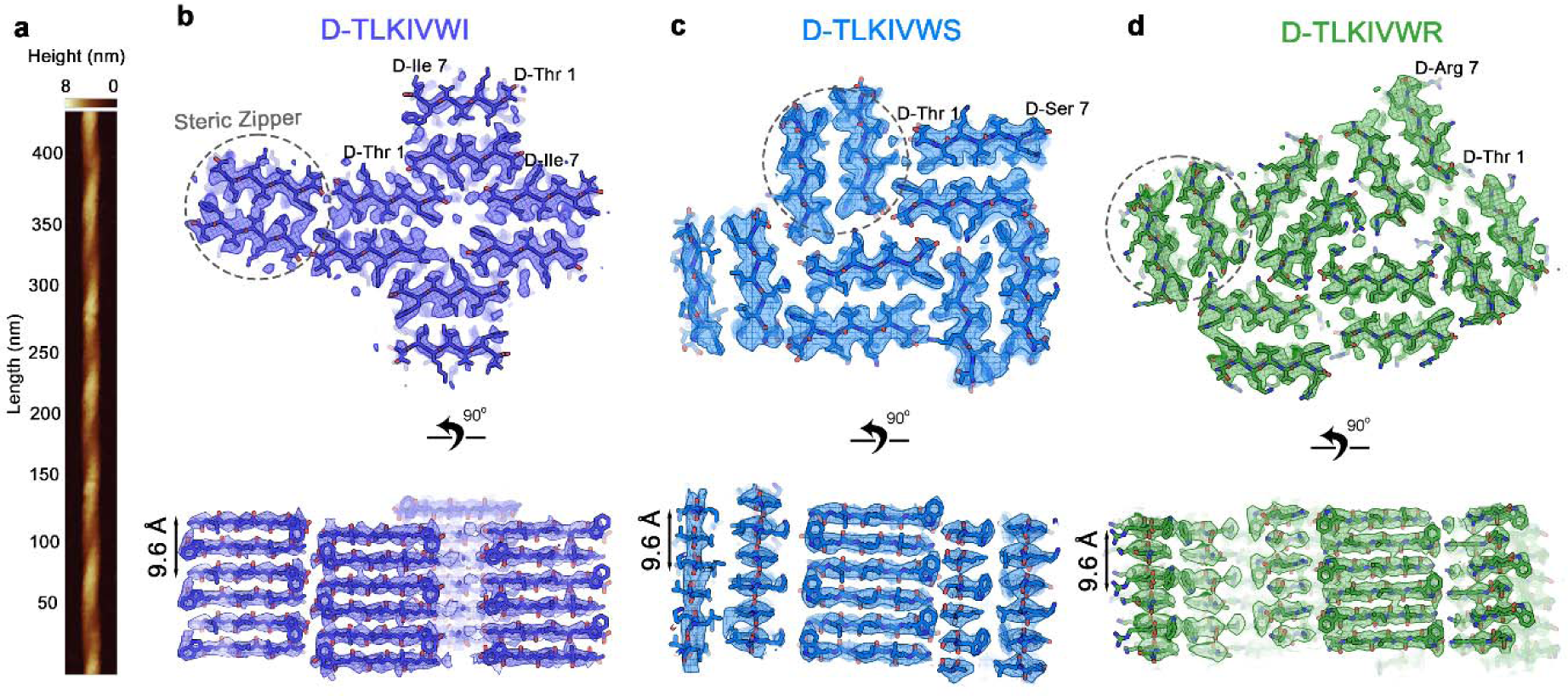
D-TLKIVWX (X = I, S, R) peptides form right-handed helical fibrils with conserved steric zipper motifs. **a**, Representative atomic force microscopy (AFM) image of D-TLKIVWI fibrils. **b-d,** Views down the fibril axes (upper row) and perpendicular to the fibril axes (lower row) of the amyloid-like fibrils formed by (**b**) D-TLKIVWI, (**c**) D-TLKIVWS, (**d**) D-TLKIVWR, as determined by cryo-EM. The conserved steric zipper motifs are represented by grey dotted ellipses.

The D-TLKIVWX (X = I, S and R) fibrils are each composed of different numbers of protofilaments and these protofilaments associate in different patterns, but all the protofilaments share the same underlying structural motif known as a “steric zipper”^28^, a pair of β-sheets mated together by an interface of snugly fitting sidechains (Fig. 3b-d and Extended Data Fig. 7a,b). In addition, the steric zippers formed by D-TLKIVWX (X = I, S and R) all share the same symmetry pattern in which antiparallel β-sheets mate together by interfacing sidechains of Leu2, Ile4, and Trp6 (an example of “class 5” symmetry ^29^). As a result, the helical rise of D-TLKIVWX (X = I, S and R) amyloid-like fibrils is 9.56 Å, instead of the 4.80 Å spacing that is common among pathogenic amyloid fibrils. Notably, the sidechain of the seventh residue faces outward from the steric zipper interface in all cases. The identity of the seventh sidechain appears to affect the geometry of association between protofilaments but does not affect the symmetry of the protofilament itself. Thus, the ability to disassemble AD-tau fibrils seems linked more strongly to the steric zipper symmetry (which is conserved among all three functional D-peptides), rather than the pattern of association between zippers (which differs among the three D-peptides).

Importantly, the structures suggest why the absence of a seventh residue in D-TLKIVW prohibits its fibril formation, and therefore presumably its inability to disassemble tau. Removal of the seventh residue would destabilize fibril formation by reducing the number of backbone hydrogen bonds (10 vs. 14) and increasing the distance between the negative charge of the C-terminal carboxylate and the compensating positive charge at the N-terminal amine of the adjacent strand (5.0 Å vs. 2.8 Å) (Extended Data Fig. 7c).

### D-TLKIVWX protofilaments grow along left-handed AD-tau fibrils

To elucidate how D-TLKIVWX might interact with AD-tau fibrils to induce their disassembly, we cryogenically trapped complexes of D-TLKIVWX (X =I, S and R) with AD-tau fibrils at an intermediate 24-h time point with a lower ratio of D-TLKIVWX to tau (estimated 100:1) in comparison with the time-course experiment in Fig. 2a (500 :1). As a negative control, we also collected images of AD-tau fibrils in the absence of D-TLKIVWX. Helical reconstructions of the AD-tau complexed with each of the three D-peptides revealed the paired helical filament (PHF) tau polymorph^30^ (Extended Data Fig. 8), and the atoms modeled into the PHF density showed no significant structural deviations from the negative control (Supplementary Table 2). However, the cryo-EM map of PHF complexed with D-TLKIVWX (X =I, S and R) revealed residual density near Val313-Thr319 of PHF, which was absent from our control (Fig. 4a-c and Extended Data Fig. 9a,b). We attribute the residual density to D-TLKIVWX for two reasons: (1) the shape of the residual density resembles one steric zipper unit of D-TLKIVWX fibrils (Fig. 4a-c and Fig. 3b-d); (2) ^1^H-^15^N-HSQC NMR experiments indicated that D-TLKIVWX (X =I and S) interacted with ^15^N,^13^C-labeled tau K18+ monomer^31^ with chemical shift perturbation mapped to Val306-Lys311^11^ and Val313-Thr319 (Fig. 4d and Extended Data Fig. 9c,d), consistent with the location of additional density next to tau PHF. Note that a rapid precipitation of D-TLKIVWX occurred upon mixing in our NMR experiments, corresponding to initial D-peptide fibril formation, but chemical shift changes of monomeric tau were observed over time when soluble fraction of D-peptides increased. Refinement of the 3D reconstruction of D-TLKIVWX (X =I, S and R) complexed with Tau PHF achieved overall resolutions of 3.1 Å, 3.1 Å and 3.5 Å, respectively. We note that the cryo-EM density map of D-TLKIVWI is stronger than that of D-TLKIVWS/R and that D-TLKIVWR is slightly further from the core of tau PHF (Fig. 4a-c and Extended Data Fig. 9b). This difference in occupancy and positioning may explain their unequal efficiency in disassembling AD-tau fibrils (Fig. 1b).

**Fig. 4:**
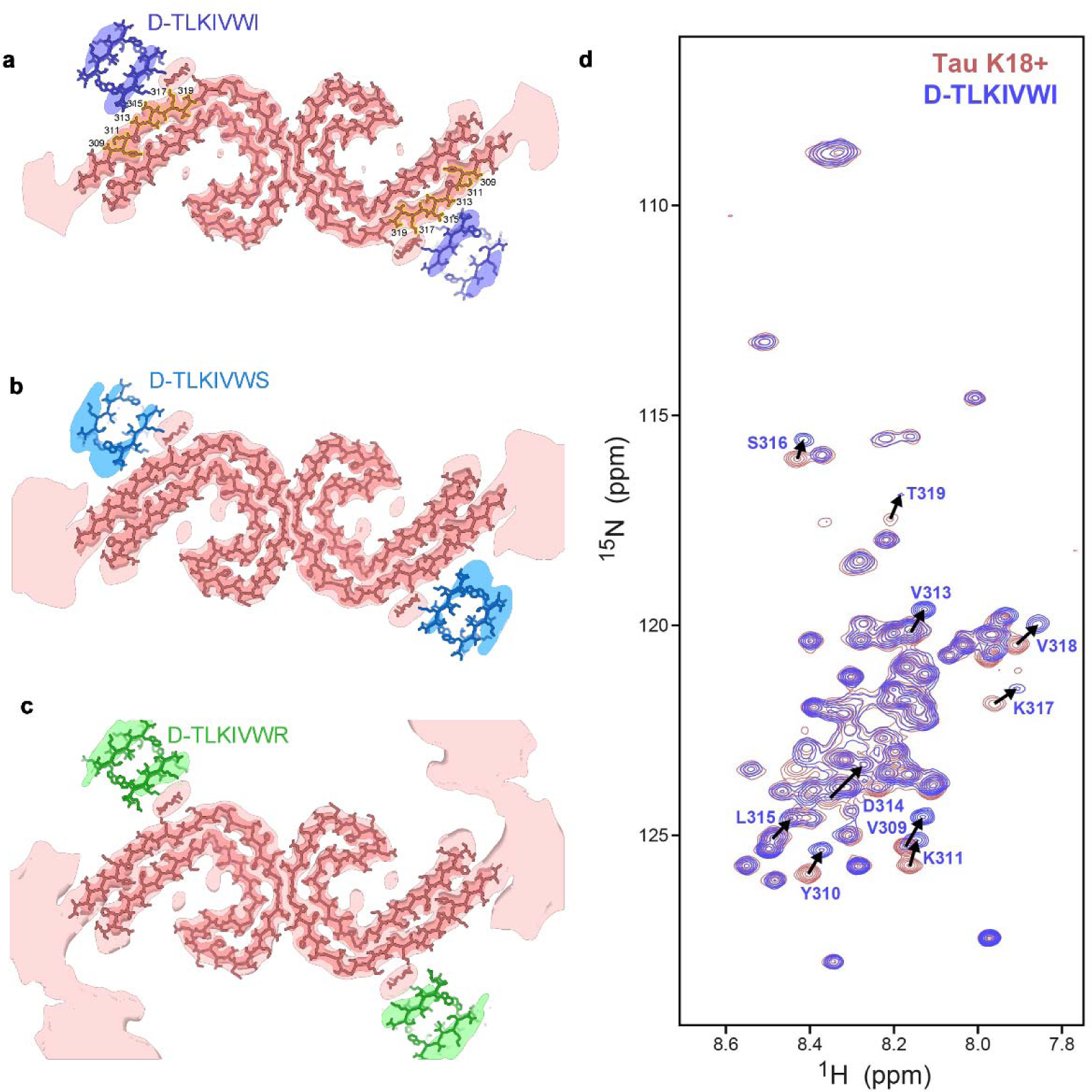
Cryo-EM structures of D-TLKIVWX (X = I, S and R) in the act of disassembling tau PHF. **a**-**c**, Cryo-EM density, and atomic models of a cross section of PHF fibrils complexed with (**a**) D-TLKIVWI, (**b**), D-TLKIVWS, and (**c**), D-TLKIVWR. Details of the PHF structures are evident in sharpened, high-resolution density maps of the PHFs (dark salmon). The D-peptides are visible in unsharpened, low-pass filtered density (7 Å) colored blue (D-TLKIVWI), marine (D-TLKIVWS), and green (D-TLKIVWR). The corresponding low-pass filtered density of the PHF is shown in light salmon. A small blob of residual density sandwiched between the PHF and each of the D-peptides was modeled with EDTA. **d**, ^1^H-^15^N HSQC spectral overlay of free ^15^N,^13^C-labeled tau K18+ (salmon), and ^15^N,^13^C-labeled tau K18+ with 10-fold molar excess of D-TLKIVWI after incubating for 2.5 months at 4 °C (blue). Representative residues with significant chemical shift changes are labeled and resonance shifts indicated with arrows. These shifted residues (Val309-Thr319) are highlighted in yellow in panel a.

## Discussion

### Strain-relief of D-TLKIVWX fibrils drives disassembly of AD-tau fibrils

Our observations lead us to propose the strain-relief mechanism of amyloid fibril disassembly illustrated in Fig. 5. As D-TLKIVWX binds to the Tau PHF, it stacks to form D-TLKIVWX amyloid-like protofilaments and these protofilaments are constrained by binding to adopt the left-handed helical twist of Tau PHF (Fig. 5a,b). Because D-TLKIVWX fibrils have an intrinsic right-handed twist (Fig. 3), the cryo-trapped structure of Figure 4a-c is metastable, and a strain develops. If not trapped by freezing, this metastable structure disassembles in hours.

**Fig. 5:**
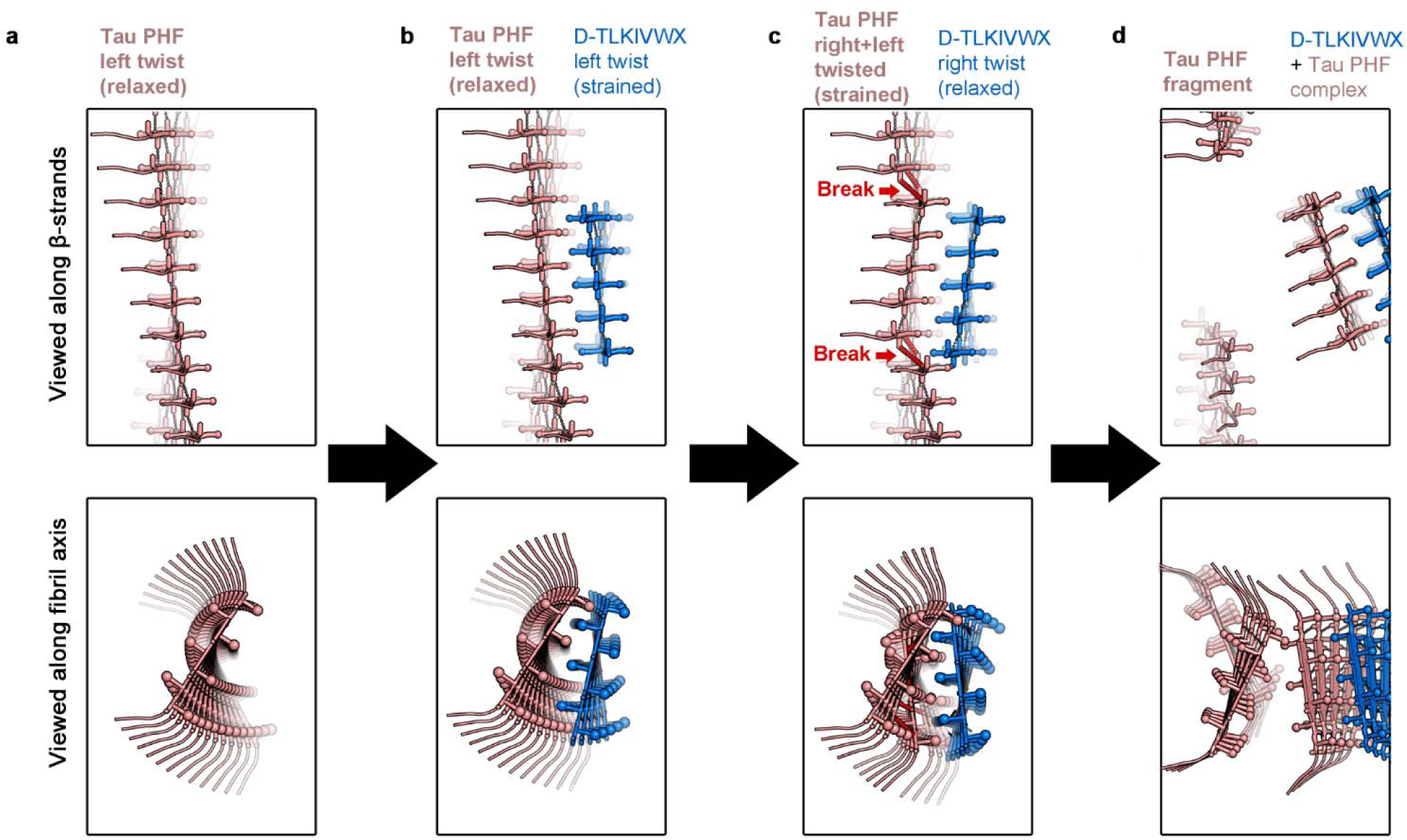
Proposed strain-relief mechanism of disassembly of tau PHF by D-TLKIVWX peptide. **a**, Diagram of left-handed tau PHF. **b**, D-TLKIVWX peptides assemble along the axis of tau PHF forming β-sheet-β-sheet interactions stabilized by steric zippers. This produces a metastable D-TLKIVWX protofilament with the left-twisted handed twist of the tau PHF. The strained left-twisted D-TLKIVWX protofilament is at a higher potential energy relative to the right-twist preferred by D-TLKIVWX peptides. **c,** Torsional strain is relieved as the D-TLKIVWX protofilament relaxes to a right-handed twist. The tau PHF sheet, in contrast, develops new strain as it struggles to maintain its interface with the now right-twisted D-TLKIVWX protofilament. Backbone hydrogen-bonds break in the tau PHF at the junction between left-and right twisted segments (red dotted lines). **d**, Solvent invasion of the tau PHF core facilitates further disassembly of the PHF. The disassembly products of the reaction are complexes of tau and D-TLKIVWX.

We propose that the strain produced by the metastable left-hand twist of D-TLKIVWX is relieved by the protofilament reversing its twist from a left-to right-handed helix (Fig. 5c). Because the D-peptide fibrils is bound to Tau PHF fibril, the reversal pulls against the Tau PHF structure. As a result, the tau residues that contact D-TLKIVWX are torn away from the native PHF contacts, thereby breaking backbone hydrogen bonds between tau molecules, and permitting solvent to invade the fibril core and further dissociate tau PHF (Fig. 5d). The disassembly products are non-seeding species (Fig. 1); they are tau-D-TLKIVWX complexes, not tau monomers (Extended Data Fig. 4c). This dynamic process is visually depicted in the Supplementary Video 1.

Our proposed strain-relief mechanism is consistent with all our experimental findings. First, D-TLKIVWX disassembles tau fibrils to which its precursor, D-TLKIVW, was designed to bind, but does not disassemble other left-handed amyloid fibrils, such as α-synuclein and wildtype hnRNPA2 LCD fibrils. Second, other analogs of D-TLKIVWX, including D-TLKIVW, D-TLKIVWP, and D-TLK(N-Me-I)VWX (X = I, S, R) fail to disassemble AD-tau fibrils, presumably because they lack the ability to form fibrils themselves. Third, the variation in efficacy of disassembly AD-tau fibrils among D-TLKIVWX can be influenced by their binding strength with AD-tau fibrils. Fourth, the concurrent disappearance of AD-tau fibrils and the newly formed D-TLKIVWX fibrils, along with the emergence of amorphous products at later time points observed in the time-course EM images (Fig. 2a) strongly suggests that the AD-tau fibrils are disassembled.

An additional observation that supports this “strain-relief” mechanism of disassembly is that L-TLKIVWX (X = C, I, S, R) all display inferior efficacy in disassembling AD-tau fibrils compared to their enantiomers D-TLKIVWX (Extended Data Fig. 10a,b). Despite exhibiting the same fibril-forming property as D-TLKIVWX^32^ (Extended Data Fig. 10c) and nearly identical binding to tau monomers (Extended Data Fig. 10d,e), the L-TLKIVWI fibrils principally possess a left-handed twist, same as tau PHF. Therefore, the L-TLKIVWX would have less structural torsion to release when they bind and assemble along the axis of tau PHF in comparison with D-TLKIVWX, resulting in their inferior efficacy in disassembling AD-tau fibrils.

The strain-relief mechanism may be a general theme of action of disruptors of amyloid fibrils. In previous work, we presented evidence that the polyphenolic compound EGCG disassembles AD-tau fibrils by stacking into a metastable amyloid-like fibril on the surface of AD-tau fibrils^9^. A subsequent change in which aromatic rings of EGCG curve into a more stable conformation could provide the energy to disassemble stable AD-tau fibrils. Thus, both disassembling actions of the very different compounds EGCG and D-TLKIVWX on AD-tau fibrils may be considered examples of strain-relief mechanisms.

In summary, we find the disassembly of AD-tau fibrils is not exclusive to D-TLKIVWC, because D-TLKIVWX (X = A, S, D, I, V, R, K, E, T) also displays this property, but with unequal efficacy. We find that the amyloid-like, fibril-forming property of D-TLKIVWX contributes to the disassembly of AD-tau fibrils. Based on cryo-EM, AFM, NMR, and other results reported here, we propose that the disassembly of AD-tau fibrils is driven by the release of torsion in D-TLKIVWX protofilaments. It remains unknown whether D-TLKIVWX disassembles tau fibrils from other tauopathies^33^. The strain-relief mechanism of amyloid disassembly proposed here may explain how diverse small molecules can provide sufficient energy to disrupt extremely stable pathogenic amyloid fibrils. This mechanism may be applied to the design of a new generation of disaggregators for tau and other pathological amyloids.

## Methods

### Chemicals and Materials

All the peptides were synthesized by GenScript and purified to ≥98%, as determined by mass spectrometry and HPLC (GenScript Corp, Piscataway, NJ).

### Recombinant protein expression and purification

Unlabeled recombinant tau K18+ (residues Gln244-Glu380 of 4R tau) was expressed in a pNG2 vector in BL21-Gold *E. coli* cells grown in LB to an A_600_ = 0.8. Cells were induced with 0.5 mM isopropyl 1-thio-β-D-galactopyranoside (IPTG) for 3 h at 37 °C and lysed by sonication in 20 mM MES buffer (pH 6.8) with 1 mM EDTA, 1 mM MgCl_2_, 1 mM dithiothreitol (DTT), and HALT protease inhibitor before the addition of NaCl (500 mM final concentration). The lysate was boiled for 20 min and then clarified by centrifugation at 15,000 rpm for 15 min and dialyzed to 20 mM MES buffer (pH 6.8) with 50 mM NaCl and 5 mM DTT. Dialyzed lysate was purified on a 5-mL HiTrap SP HP ion exchange column and eluted over a gradient of NaCl from 50 to 550 mM. Protein was further purified on a HiLoad 16/600 Superdex 75 pg column (GE Healthcare) in 10 mM Tris (pH 7.6) with 100 mM NaCl and 1 mM DTT and concentrated to 20-60 mg/mL by ultrafiltration using a 3-kDa cutoff filter (Millipore-Sigma, Burlington, MA).

Isotopically labeled tau K18+ proteins for NMR were grown in M9 H_2_O media supplemented with ^15^NH_4_Cl (and ^13^C-glucose) as the sole nitrogen (and carbon) source. Protein expression was induced with 1 mM IPTG at 37 °C for 6 hours. The purification was same as for the unlabeled protein.

The construct for overexpression of mCherry-hnRNPA2-LCD fusion protein was provided by Dr. Masato Kato of University of Texas, Southwestern. Protein overexpression and purification procedures followed the same protocol reported previously^34^.

### Extraction of tau fibrils from AD patient brains

Frozen brain tissues were weighed, diced into small pieces, and resuspended in 10 mL/gram of sucrose buffer (800LJmM NaCl, 10% sucrose, 10 mM Tris-HCl, pH 7.4, 0.1 mM EDTA, 1 mM DTT) supplemented with 1:100 (v/v) Halt protease inhibitor (Thermo Scientific). Resuspended tissue was homogenized using a Polytron homogenizer (Thomas Scientific) and centrifuged at 20,000 × g at 4 °C for 20 minutes. The crude supernatant was treated with N-lauroylsarcosinate (1% [w/v] final concentration) and shaken at room temperature (22 °C) for 1 h. The supernatant was then centrifuged at 100,000 × g for 1 h at 4 °C. The sarkosyl-insoluble pellet was resuspended in washing buffer (10LJmM Tris–HCl, pH 7.4, 800LJmM NaCl, 5LJmM EDTA, 1 mM EGTA, 1LJmM DTT, 10% sucrose) and centrifuged at 20,100 x g for 30LJmins at 4 °C. After centrifugation, the supernatant was further centrifuged at 100,000 × g (Beckman Coulter, Optima MAX-XP) for 1 h at 4 °C. Finally, the purified sarkosyl-insoluble pellet was resuspended in 250 μl 20LJmM Tris-HCl, pH 7.4, 100LJmM NaCl and stored at -80 °C.

### Thioflavin T (ThT) assay

Kinetic fluorescence data were collected in a microplate reader (FLUOstar Omega, BMG Labtech) at 37 °C with double orbital shaking at 700 rpm. Fluorescence measurements were recorded every 10 mins with excitation and emission wavelengths of 440 and 480 nm. All samples were added in triplicate and experiments were repeated at least twice.

### Dot blot assay

Purified AD-tau fibrils from brain extract were incubated with 500 μM L/D-TLKIVW, D-TLKIVWX (X = C, A, S, D, I, V, R, K, E, P, T), D-TLK(N-Me-I)VWX (X =I, S and R), L-TLKIVWX (X = C, I, S and R) at 37 LJ for 48 h, respectively. 2.5 μL of samples were added on nitrocellulose membrane (0.2 µm, Bio-Rad). The membrane was blocked by 5% (w/v) nonfat dry milk in TBS-T (T = 0.1% (v/v) Tween-20) at room temperature for 1 hr. After blocking, the membrane was incubated with GT38 antibody obtained from Virginia Lee’s lab (1:1000) in 5% (w/v) milk in TBS-T at 4 LJ overnight. Then, the membrane was washed in TBS-T three times for 5 minutes each and incubated with goat anti-mouse IgG HRP (1:5000, cat# AB205719, Abcam) in TBS-T for 1 h at room temperature. The membrane was washed three more times, and the signal was developed with Pierce^TM^ ECL western blotting substrate (170-5061, BioRad).

### Negative stain transmission electron microscopy (TEM)

6 μL of sample was applied to a glow discharged carbon coated electron microscopy grid (CF150-Cu, Electron Microscopy Sciences) for 5 minutes. Then grids were stained with 2% uranyl acetate for 2 minutes. Samples were visualized using a FEI Tecnai T12 Quick room temperature transmission electron microscope equipped with a Gatan 2,048 x 2,048 CCD camera operated at an acceleration voltage of 120 kV.

### AD brain tau fibril seeding in tau biosensor cell line

HEK293 cell lines stably expressing tau-K18-YFP were engineered by Marc Diamond’s laboratory at the University of Texas Southwestern Medical Center and used without further characterization or authentication. Cells were maintained in Dulbecco’s modified Eagle’s medium (DMEM, Life Technologies, cat# 11965092) supplemented with 10% (v/v) fetal bovine serum (Life Technologies, cat# A3160401), 1% antibiotic-antimycotic (Life Technologies, cat# 15140122), and 1% Glutamax (Life Technologies, cat# 35050061) at 37 °C, 5% CO_2_ in a humidified incubator. AD-tau fibrils were incubated with D-TLKIVWX (X = A, S, D, I, V, R, K, E, P, T) (2.5, 5, 10, 20, 50, 75, 100 μM) at 4 °C overnight and sonicated in a cup horn water bath for 3 min. Then these disassembly products of AD-tau were mixed with 1 volume of Lipofectamine 3000 prepared by diluting 1μL of Lipofectamine 3000 (Life Technologies, cat# 2729899) in 19 μL Opti-MEM (Life Technologies, cat# 31985070). After 20 min, 10 μL of fibrils were added to 90 μL tau biosensor cells. After 24 hours of incubation, the number of seeded aggregates was determined by imaging the entire well of a 96-well plate in triplicate using a Celigo image cytometer (Nexcelom) in the YFP channel. The data analysis was described before. For high-quality images, cells were photographed on a ZEISS Axio Observer D1 fluorescence microscope using the EGFP fluorescence channel.

### Cell toxicity of AD-tau disassembly species

AD-tau fibril disassembly species were produced by incubating AD-tau fibrils (estimated 1 μM) with D-TLKIVWX (5, 10, 20, 50, or 100 μM) at 4 °C overnight. Neuro 2A (N2a) cells were cultured in DMEM supplemented with 10% (v/v) fetal bovine serum, 1% antibiotic-antimycotic, and 1% Glutamax in a 5% CO_2_ humidified environment at 37 °C. Cells were plated at a density of roughly 6,000 cells/well on 96-well plates in 90 μL of fresh medium. After 24 h, 10 μL of the above AD-tau fibril disassembly species were added and the cells were incubated for another 24 h at 37 °C. Cytotoxicity was measured utilizing an MTT assay.

### Western blot

Purified brain-extracted AD-tau fibrils were incubated with 500 μM D-TLKIVWI at 37 LJ for 48 h. The sample was centrifuged at 21,000 x g for 30 min at 4 LJ (Eppendorf Centrifuge 5424R). Western blot was performed with anti-tau rabbit polyclonal antibody (1:1000, Dako A0024) and anti-rabbit HRP-conjugated secondary antibody (1:5000, Thermo Fisher Scientific).

### X-ray diffraction (XRD)

D-TLKIVWX (X = I, S, R) peptides were dissolved to 10 mM in deionized water and incubated at room temperature quiescently for three days. Peptide fibrils were aligned by pipetting the suspension in a 3 mm gap between two fire-polished glass rods and drying overnight. The aligned fibrils were cooled to 100 K. Diffraction data was collected on a FR-E+ rotating anode x-ray generator (Rigaku) equipped with a R-AXIS HTC imaging plate detector (Rigaku). Cu K-α x-ray beam with 1.5406 Å wavelength was used, and the detector was placed 78 mm from the sample. Diffraction images were visualized using ADXV (The Scripps Research Institute).

### Atomic force microscopy (AFM)

4 mM D-TLKIVWI in deionized water was shaken at room temperature for three days and diluted into distilled water in a 1:10 ratio. Then, 5 μL of diluted sample was deposited onto freshly cleaved mica and incubated for 10 min. The sample was rinsed with Milli-Q water and dried under a stream of nitrogen gas. AFM images were collected using a Dimension Icon microscope (Bruker) in PeakForce Tapping mode using ScanAsyst-HR probes. Each collected image had a scan size of 3 x 3 μm and 2048 x 2048 pixels and was collected using a scan rate of 0.494 Hz. Nanoscope Analysis software (Version 2.0, Bruker) was used to process the image data by flattening the height topology data to remove tilt and scanner bow. Fibrils were traced and computationally straightened from collected AFM images in Matlab using Trace_y^35^.

### Cryo-EM samples

D-TLKIVWI fibrils were optimized by shaking 4 mM D-TLKIVWI in deionized water at room temperature for three days. 10 mM D-TLKIVWS/R in deionized water formed fibrils when left undisturbed for three days at room temperature. For D-TLKIVWR fibrils, the pH of the peptide solution was adjusted to 7.0. Prior to cryo-EM grid preparation, AD-tau fibrils in a buffer comprised of 20 mM Tris-HCl pH 7.4, 100 mM NaCl were pre-incubated at 37 °C with final concentration of 100 µM D-TLKIVWX (X = I, S, R) from 10 mM stocking solution in water for 24 hours. Control tau fibrils from the same brain donor were treated identically except for the addition of D-TLKIVWX.

### Cryo-EM data collection and processing

To prepare the cryo-EM grids, we applied 2.5 μl of sample solution onto Quantifoil 1.2/1.3 200 mesh electron microscope grids glow-discharged for 2 minutes in a Pelco easiGlow unit before use. Grids were plunge-frozen into liquid nitrogen-cooled liquid ethane inside a Vitrobot Mark IV (FEI) vitrification robot after blotting. Cryo-EM data of D-TLKIVWR and D-TLKIVWS fibrils were collected on a Titan Krios transmission electron microscope (Thermo Fisher Scientific) located at the National Center for Cryo-EM Access and Training, which is equipped with a Bioquantum/K3 direct detection camera (Gatan), operated with 300 kV acceleration voltage and an energy slit width of 20 eV, automated with Leginon software package^36^. Super-resolution movies were collected with a calibrated pixel size of 1.067 Å/pixel (0.5335 Å/pixel in super-resolution movie frames) and a dose per frame of ∼1.5 e-/Å^2^. A total of 40 frames with a frame rate of 12 Hz were taken for each movie, resulting in a final dose of ∼60 e-/Å^2^ per image. D-TLKIVWI fibrils were collected on a Titan Krios located at the HHMI Janelia Research Campus, which is equipped with a cold-FEG source (CFEG), a Selectris X energy filter and a Falcon 4i direct detection camera (TFS), operated with 300 kV acceleration voltage and an energy slit width of 6 eV, and automated with the SerialEM software package^37^. Electron Event Representation (EER) files were collected with a calibrated pixel size of 0.94 Å/pixel and a dose per raw frame of 0.0244 e-/Å^2^, resulting in 55 e-/Å^2^ per image. AD-tau/D-TLKIVWX (X = I, S and R) were collected similarly as D-TLKIVWI fibrils, although manually targeted in SerialEM package. The AD-tau control was collected on a Titan Krios/Bioquantum/K3 setup located at the Stanford-SLAC Cryo-EM Center, operated with 300 kV acceleration voltage and an energy slit width of 20 eV, automated with EPU (TFS). (See details in Supplementary Table 1,2).

Movies and EER files were motion-corrected in RELION^38^ and binned to pixel sizes according to Supplementary Table 1,2. CTF estimation was performed using CTFFIND4^39^. AD-tau/D-TLKIVWX fibrils were manually picked using e2helixboxer.py from EMAN2^40^. D-TLKIVWX fibrils and AD-tau control particle picking was initially done manually using e2helixboxer.py from EMAN2 for about 100 images as a training set for crYOLO^41^. CrYOLO was then trained with default parameters and was used to pick the rest of the images. Particle extraction, two-dimensional classification, three-dimensional classification, and 3D refinement were performed in RELION^42^. Briefly, particles were initially extracted using a larger box size of 640 pixels with two-fold binning. 2D classification was then performed with all particles to eliminate bad particles and group particles into polymorphs if necessary. Particles from each polymorph were selected, extracted with smaller box sizes at native pixel sizes of detectors (binning=1), further “purified” using 2D classification and subjected to 3D classification, which was done initially with one class and then with three classes, using a Gaussian cylinder as the initial model. The best 3D classes were selected, and corresponding particles were finally refined with 3D auto-refine for the reported maps. (See details in Table Supplementary Table 1,2). Part of the Cryo-EM data processing used Expanse GPU at San Diego Supercomputer Center through allocation BIO230174 from the Advanced Cyberinfrastructure Coordination Ecosystem^43^.

### Atomic model building

Our starting atomic model of D-TLKIVWI was an ideal β-strand. It was manually adjusted to fit the electrostatic potential map using Coot^44^ and automated refinement was performed using Phenix^45^. To facilitate good rotamer geometry, the initial building and refinement was performed using a map with handedness chosen so that the amino acid residues appeared to be levorotary rather than dextrorotary. In this way, we could take advantage of the rotamer library in Coot which exists for L-amino acids, but not for D-amino acids. In the final step of refinement, the map and coordinates were inverted to the correct hand, consistent with D-amino acids. The starting models for D-TLKIVWS and D-TLKIVWR were adapted from the refined D-TLKIVWI structure. All atomic models were refined in successive rounds using Coot for manual building and Phenix for automated refinement. Model validation statistics of all three D-peptide structures are reported in Supplementary Table 1.

Our starting atomic model of the complex between AD-tau PHF and D-TLKIVWI was built by manually orienting coordinates of the tau PHF (PDB ID 7nrv)^46^ to fit the electrostatic potential map using Coot^44^ and then refined with Phenix. Coordinates of a pair of β-sheets were extracted from the D-TLKIVWI structure described above and manually docked on the surface of the PHF using guidance from the 3.1 Å cryoEM map, as well as the low-pass filtered map (7 Å). We noted a blob of residual density situated at the end of three lysine side chains: K317 and K321 of tau and K3 of the D-peptide. Whatever molecule produced this residual density does not depend on the presence of the D-peptide to bind to tau, since a similar blob was evident in our PHF negative control lacking D-peptide. Indeed, the presence of this residual density was noted in the original structure report of AD-tau PHFs^30^, and even noted in maps from PHFs produced with recombinant tau (PDB ID 7ql4)^47^. The chemical environment of this blob suggests that the blob originates from an anion, but the density is not sufficiently detailed to uniquely identify the chemical species. It is roughly the size of a pair of phosphate ions. We chose to model this residual density with ethylenediaminetetraacetate (EDTA) because it fits the density, caries the expected negative charges to complement the positive charge on K317 and K321, and we know that EDTA was included in the buffer used for PHF purification. The starting models for tau complexed with D-TLKIVWS and D-TLKIVWR were obtained using an analogous procedure. The final refined coordinates for these two complexes do not include the D-peptides because density for the peptides was visible only in the low-pass filtered maps, and not in the high-resolution map (Figure 4 and Extended data Figure 9a).

### NMR spectroscopy

NMR samples were ∼0.5 mL of 0.1 mM ^15^N,^13^C-labeled tau K18+ protein in 100 mM KCl, 20 mM NaH_2_PO_4_, 1 mM TCEP, 5%/95% D_2_O/H_2_O, pH 7.0 without or with 1 mM D-TLKIVWI, D-TLKIVWS, L-TLKIVWI. All NMR spectra were acquired at 25 °C with Bruker Avance III HD 600 MHz spectrometer equipped with QCI HCNP cryoprobe or Avance Neo 800 MHz spectrometer equipped with TCI HCN cryoprobe. Backbone assignments for both free tau K18+ and D-TLKIVWI bound tau K18+ were carried out using HNCACB, CBCA(CO)NH and C(CO)NH NMR experiments. NMR spectra were acquired with Topspin (Bruker), processed with NMRPipe^48^, and analyzed with NMRFAM-Sparky^49^.

For the chemical shift perturbation (CSP) analysis, the overall change in chemical shift Δ was calculated between the free and bound states of tau K18+ protein as^50^:

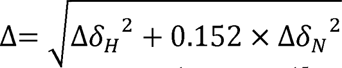

where Δδ_H_ and Δδ_N_ are the differences between the ^1^H_N_ and ^15^N chemical shifts of the two states being compared.

### Statistical analysis

Graphs are expressed as means + standard deviation (SD) and data were analyzed using SPSS 25 statistical analysis software (SPSS, Chicago, IL, USA). The one-way analysis of variance (ANOVA) was used to analyze difference among multiple groups. Statistical differences for all tests were considered significant at the \**p* < 0.05, \*\**p* < 0.01, \*\*\**p* < 0.001 levels, NS, not significant.

### Reporting summary

Further information on research design is available in the Nature Research Reporting Summary linked to this paper.

## Data availability

Cryo-EM maps and atomic models of the fibrils from peptides D-TLKIVWX (X = I, S, R) alone have been deposited in the Electron Microscopy Data Bank (EMDB) and PDB under the accession codes EMD-44181/9B4I (D-TLKIVWI), EMD-44182/9B4J (D-TLKIVWS), and EMD-44183/9B4K (D-TLKIVWR). Cryo-EM maps and atomic models of paired helical filament of Tau from Alzheimer’s patient in complex with the three peptides have been deposited in the same databases under the accession codes EMD-44184/9B4L (Tau-D-TLKIVWI), EMD-44185/9B4M (Tau-D-TLKIVWS), and EMD-44186/9B4N (Tau-D-TLKIVWR). Cryo-EM map and atomic model of the paired helical filament from the same patient have been deposited under the accession codes EMD-44187/9B4O. All data presented in this article are available within the figures and Supplementary Information files. All other data are available from the corresponding authors upon reasonable request.

## Supporting information

Cryo-EM statistics of D-TLKIVWX (X = I, S, R) fibrils

Cryo-EM statistics of AD-tau and AD-tau-D-TLKIVWX (X = I, S, R) complex

Strain-relief of D-TLKIVWX fibrils drives disassembly of AD-tau fibrils

## Abbreviations

Aβ: amyloid beta protein
AD: Alzheimer’s disease, AD-tau, tau fibrils extracted from AD brains
All D-peptides TLKIVWX (D-TLKIVWX): in which X can be any enantiomorph of the 20 coded amino acid residues
PHFs: paired helical filaments, the most abundant polymorph of AD-tau fibril

## Acknowledgments

We thank the donors and their families for the AD brain tissue; without whom this work would not have been possible. We acknowledge NIH 1R01AG070895 (D.S.E.), NIH RF1AG065407 (D.S.E.), DOE-FC02-02ERG and Alzheimer’s Association Research Fellowship AARF-21-848751 (K.H.) for support. The authors thank Dr. Marc Diamond for sharing the YFP-labeled tau K18 expressing HEK293 cells. We thank Virginia Lee for generously gifting the GT38 antibody. Some of this work was performed at the National Center for CryoEM Access and Training (NCCAT) and the Simons Electron Microscopy Center located at the New York Structural Biology Center, supported by the NIH Common Fund Transformative High Resolution Cryo-Electron Microscopy program (U24 GM129539), and by grants from the Simons Foundation (SF349247) and NY State Assembly. We also thank the staff at the HHMI Janelia CryoEM Facility for help and support. Some of this work was performed at the Stanford-SLAC Cryo-EM Center (S2C2) supported by the NIH Common Fund Transformative High Resolution Cryo-Electron Microscopy program (U24 GM129541). This work used Expanse GPU at San Diego Supercomputer Center through allocation BIO230174 from the Advanced Cyberinfrastructure Coordination Ecosystem: Services & Support (ACCESS) program, which is supported by National Science Foundation grants #2138259, #2138286, #2138307, #2137603, and #2138296. We acknowledge support of NMR equipment by grants from NIH (S10OD016336 and S10OD025073) and from DOE (DE-FC02-02ER63421). Y.Y. was supported in part by R35GM131901 to J.F.

## Author contributions

The project was conceived and designed by K.H. and D.S.E. Cell seeding experiment was performed by J.L.D. and K.H.. Dot blot, TEM and western blot were performed by K.H. and J.P.. J.Z. and J.L.D. provided the recombinant tau K18+. J.P. extracted tau fibrils from AD patient brains with the help of X.C.. X-ray diffraction was done by K.H. and Y.X.J.. AFM was carried out by L.L.. Cryo-EM grids were prepared by K.H., Y.X.J. and P.G.. Cryo-EM data were collected by P.G. with assistance from S.Y., Z.Y., Y.X.J. and D.R.B.. K.H. and P.G. processed cryo-EM data with assistance from Y.X.J., D.R.B., X.C. and J.L.. M.R.S. and K.H. built atomic models. Isotopically labeled tau K18+ was purified by K.H. NMR experiments were conducted and analyzed by Y.Y. and J.F.. The manuscript was prepared by K.H., M.R.S., P.G., and D.S.E. with contributions from all the other authors.

## Additional information (containing supplementary information line (if any) and corresponding author line)

**Supplementary Table 1.** Cryo-EM statistics of D-TLKIVWX (X = I, S, R) fibrils

**Supplementary Table 2.** Cryo-EM statistics of AD-tau and AD-tau-D-TLKIVWX (X = I, S, R) complex

**Supplementary Video 1.** Strain-relief of D-TLKIVWX fibrils drives disassembly of AD-tau fibrils. The structure-based Strain-Relief hypothesis posits that the interaction of right-handed-twisting fibrils formed by peptide D-TLKIVWI with left-handed-twisting fibrils of protein AD-tau produces strain in both fibrils and strain is relieved by disassembly of the fibrils.

## Extended data figure legends

**Extended Data Fig. 1.**
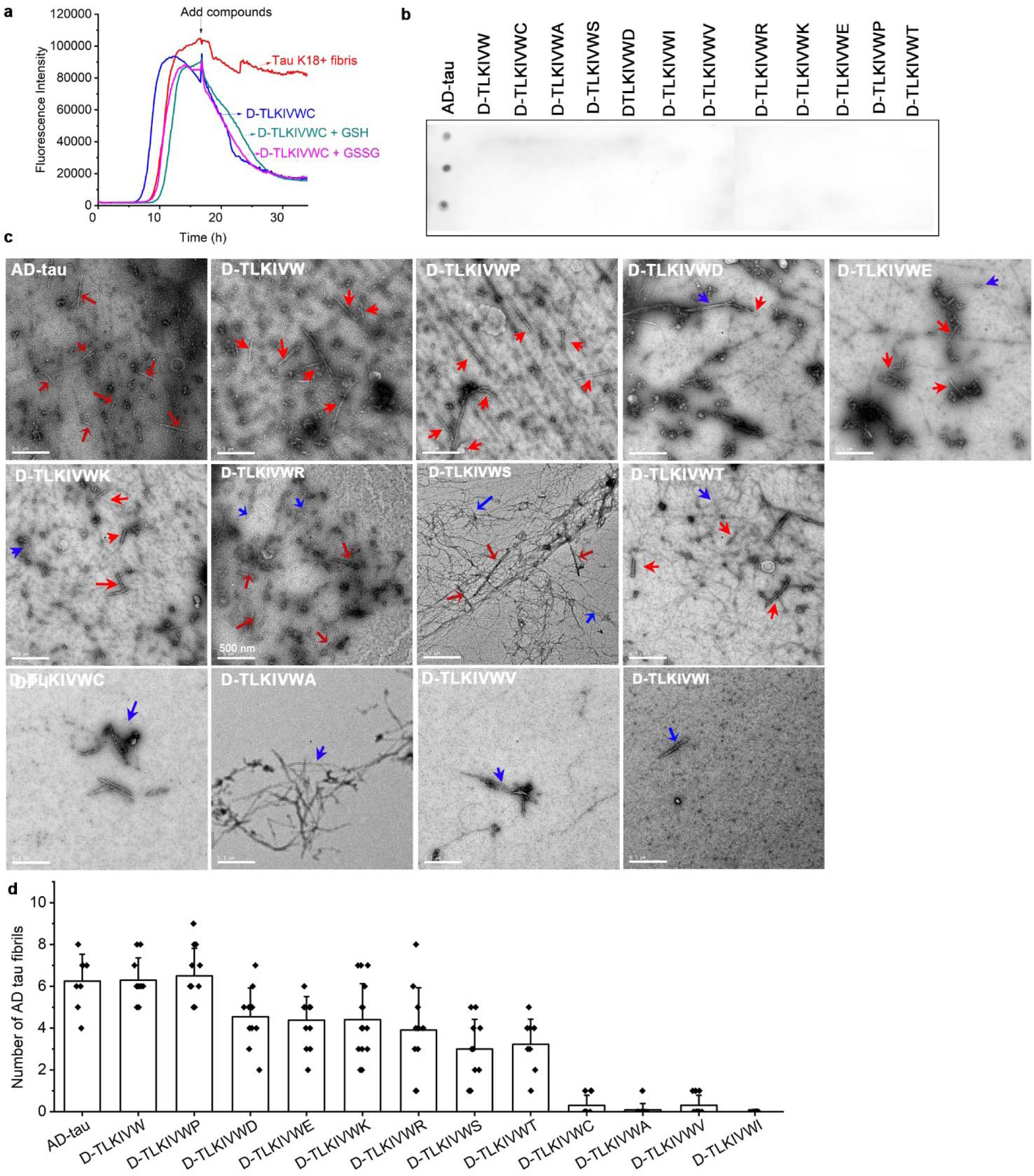
Variations of the sequence D-TLKIVWX (X = A, C, S, D, I, V, R, K, E, P, T) show varying efficacy in disassembling AD-tau fibrils. **a**, Thioflavin T (ThT) assay of 20 μM tau K18+ fibrils added with H_2_O, 50 μM D-TLKIVWC, a mix of 50 μM D-TLKIVWC and 100 μM glutathione (GSH), or a mix of 50 μM D-TLKIVWC and 100 μM Glutathione disulfide (GSSG), respectively. **b**, Dot blot staining of D-TLKIVW, D-TLKIVWX (X= C, A, S, D, I, V, R, K, E, P, T) alone using the GT38 antibody. AD-tau fibril was stained as a positive control. **c**, Representative TEM images of AD-tau fibrils after incubation with 500 μM D-TLKIVW or D-TLKIVWX (X= C, A, D, V, K, E, P, T) at 37 LJ for 48 h. The AD-tau fibrils and newly formed fibrils are labeled by red and blue arrows, respectively. Scale bars, 500 nm. **d**, Quantification of AD-tau fibrils in panels c. (*n* = 10 TEM images).

**Extended Data Fig. 2.**
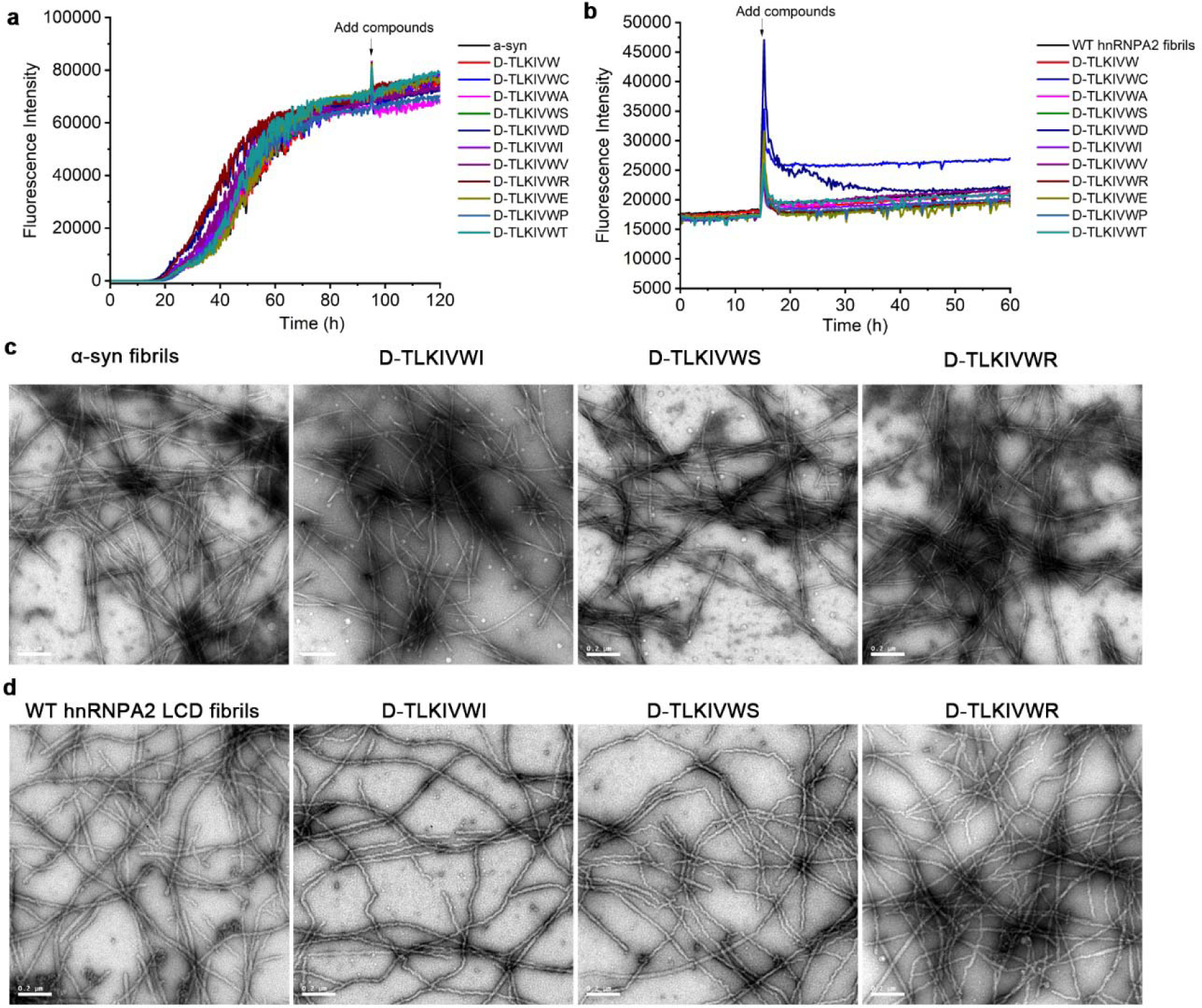
D-TLKIVWX does not disassemble α-syn fibrils or wild-type (WT) hnRNPA2 low complexity domain (LCD) fibrils. **a**, ThT assay of 25 μM α-syn fibrils incubated with 50 μM fresh D-TLKIVW or D-TLKIVWX (X = C, A, S, D, I, V, R, E, P, T) at 95 h time-point. **b**, ThT assay of 10 μM WT hnRNPA2 low complexity domain (LCD) fibrils incubated with 100 μM fresh D-TLKIVW or D-TLKIVWX (X = C, A, S, D, I, V, R, E, P, T) at 15 h time-point. **c**, Representative TEM images of a-syn fibrils alone and after incubation with 50 μM D-TLKIVWX (X = I, S and R) for 24 hours. **d**, Representative TEM images of WT hnRNPA2 LCD fibrils alone and after incubation with 100 μM D-TLKIVWX (X = I, S and R) for 55 hours. Scale bars, 0.2 µm.

**Extended Data Fig. 3.**
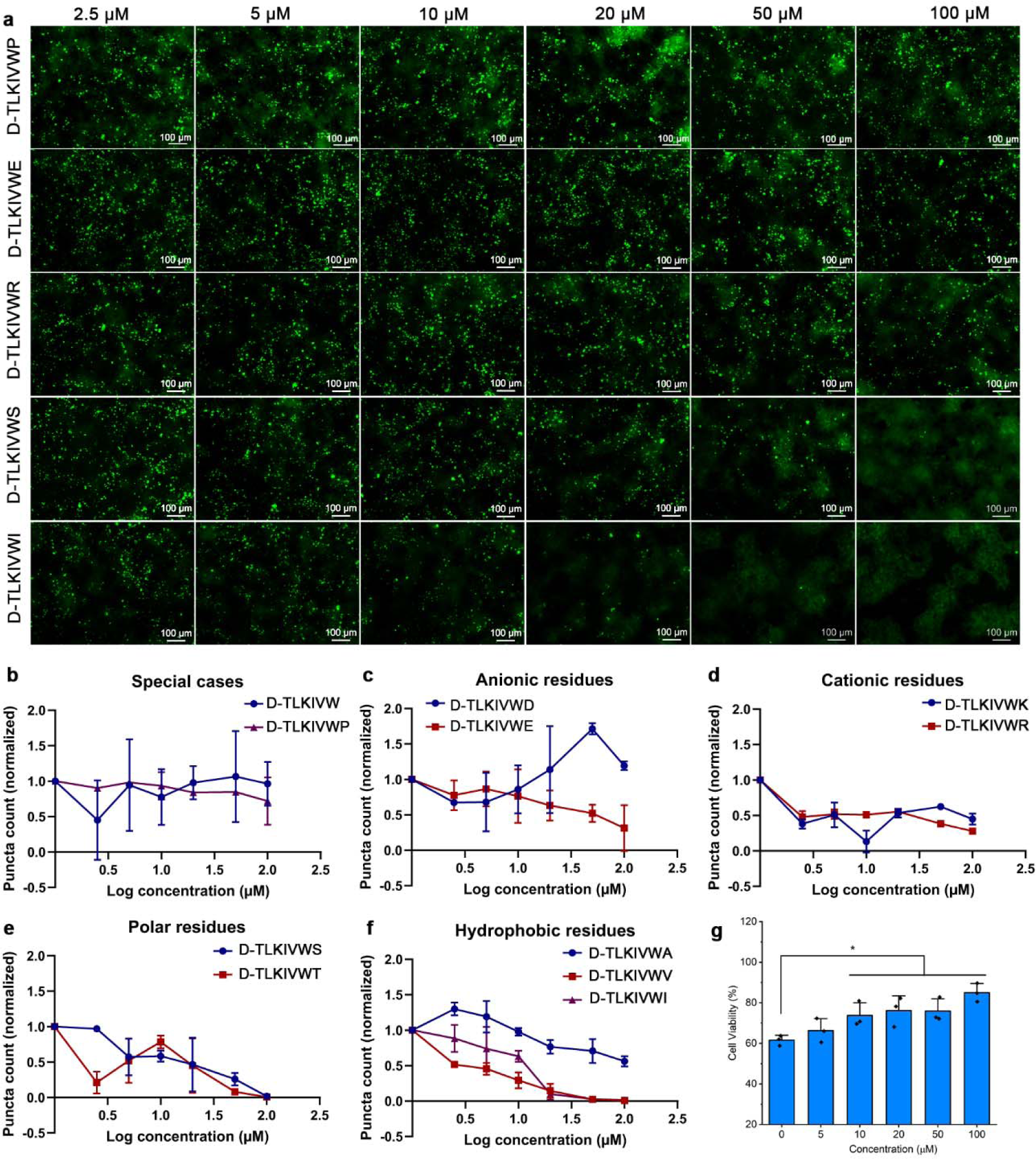
AD-tau disassembly products are non-seeding species in tau biosensor cells. **a**, Representative fluorescent images of HEK293 cells expressing YFP-labeled tau K18 transfected with AD-tau fibril seeds after overnight disassembly with various concentrations of D-TLKIVWX (X = P, E, R, S, I). Scale bars, 100 μm. **b**, Quantification of puncta formed in tau biosensor cells after overnight disassembly with various concentrations of special cases D-TLKIVWP and D-TLKIVW. **c-f**, Quantification of puncta formed in tau biosensor cells after overnight disassembly with various concentrations of (**c**) D-TLKIVWX (X = D, E), (**d**) D-TLKIVWX (X = K, R), (**e**) D-TLKIVWX (X = S, T), (**f**) D-TLKIVWX (X = A, V, I). **g**, Cell viability of N2a cells treated with AD-tau fibrils that incubated overnight with D-TLKIVWS, measured by MTT assay. Results shown as Mean + SD of triplicate wells. Statistical significance was analyzed by one-way ANOVA (\**p* < 0.05).

**Extended Data Fig. 4.**
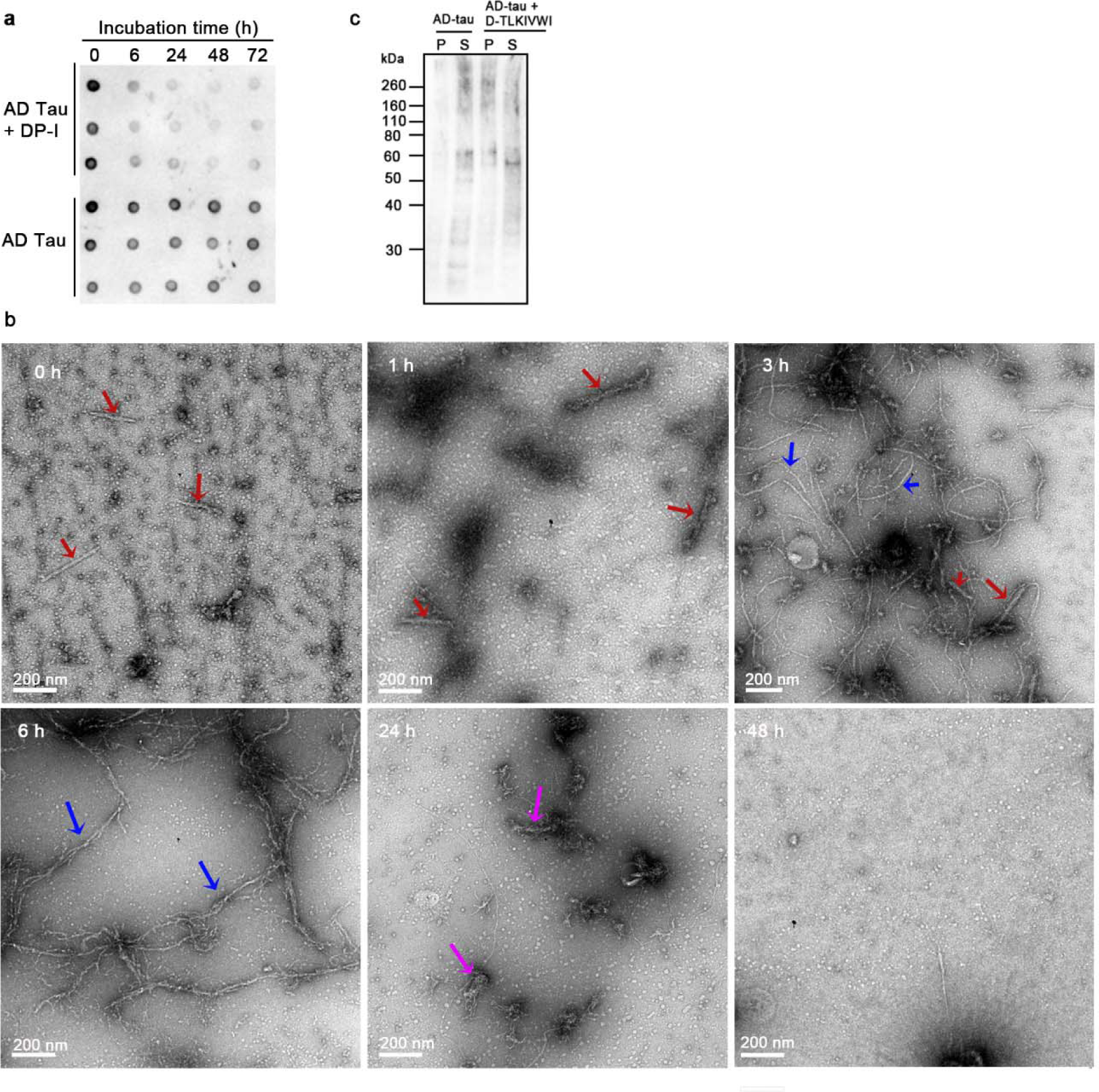
Emergence of new fibrils accompany with the disappearance of AD-tau fibrils. **a**, Time-course dot blot staining of AD-tau fibrils incubated with 500 μM D-TLKIVWI, probed by GT38 antibody. **b**, Negative stain electron micrographs of AD-tau fibrils incubated with 500 μM D-TLKIVWI at 0, 1, 3, 6, 24 and 48 h timepoints. AD-tau fibrils, newly formed D-TLKIVWI fibrils, and amorphous products are labeled by red, blue, and magenta arrows, respectively. **c**, Western blot of the supernatant and pellet of AD-tau fibrils before and after incubation with D-TLKIVWI for 48 h. The Dako antibody used here recognizes a linear tau epitope and so is unspecific for conformation. P: pellet, S: supernatant.

**Extended Data Fig. 5.**
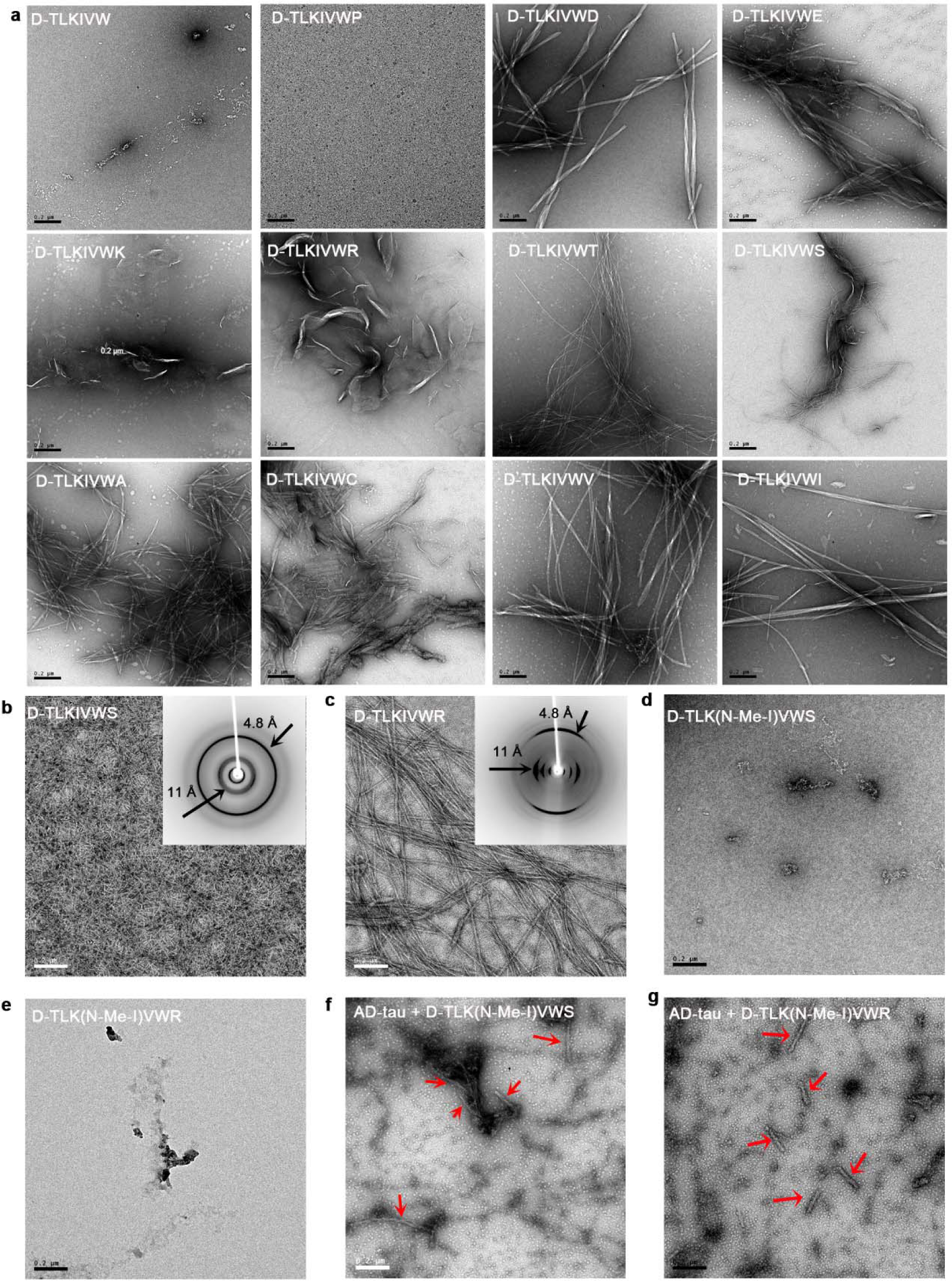
Aggregate ability of D-TLKIVWX is essential to the disassembly of AD-tau fibrils. **a,** TEM images of 500 μM D-TLKIVW, and 500 μM D-TLKIVWX (X = P, D, E, K, R, T, S, A, C, V, I) after incubation in 20 mM Tris-HCl, pH 7.4, 100 mM NaCl for 48 hours. Scale bars, 0.2 μm. **b-c**, Negative stain TEM and X-ray diffraction pattern (insets) of 10 mM (**b**) D-TLKIVWS, (**c**) D-TLKIVWR after left static in deionized water at room temperature for three days. **d-e**, Negative stain TEM image of 500 μM (**d**) D-TLK(N-Me-I)VWS, (**e**) D-TLK(N-Me-I)VWR in 20 mM Tris-HCl, pH 7.4, 100 mM NaCl at 37 LJ for 48 hours. **f-g**, Representative TEM images of AD-tau fibrils after incubation with 500 μM (**f**) D-TLK(N-Me-I)VWS, (**g**) D-TLK(N-Me-I)VWR at 37 LJ for 48 hours. AD-tau fibrils are labeled by red arrows. Scale bars, 0.2 μm.

**Extended Data Fig. 6.**
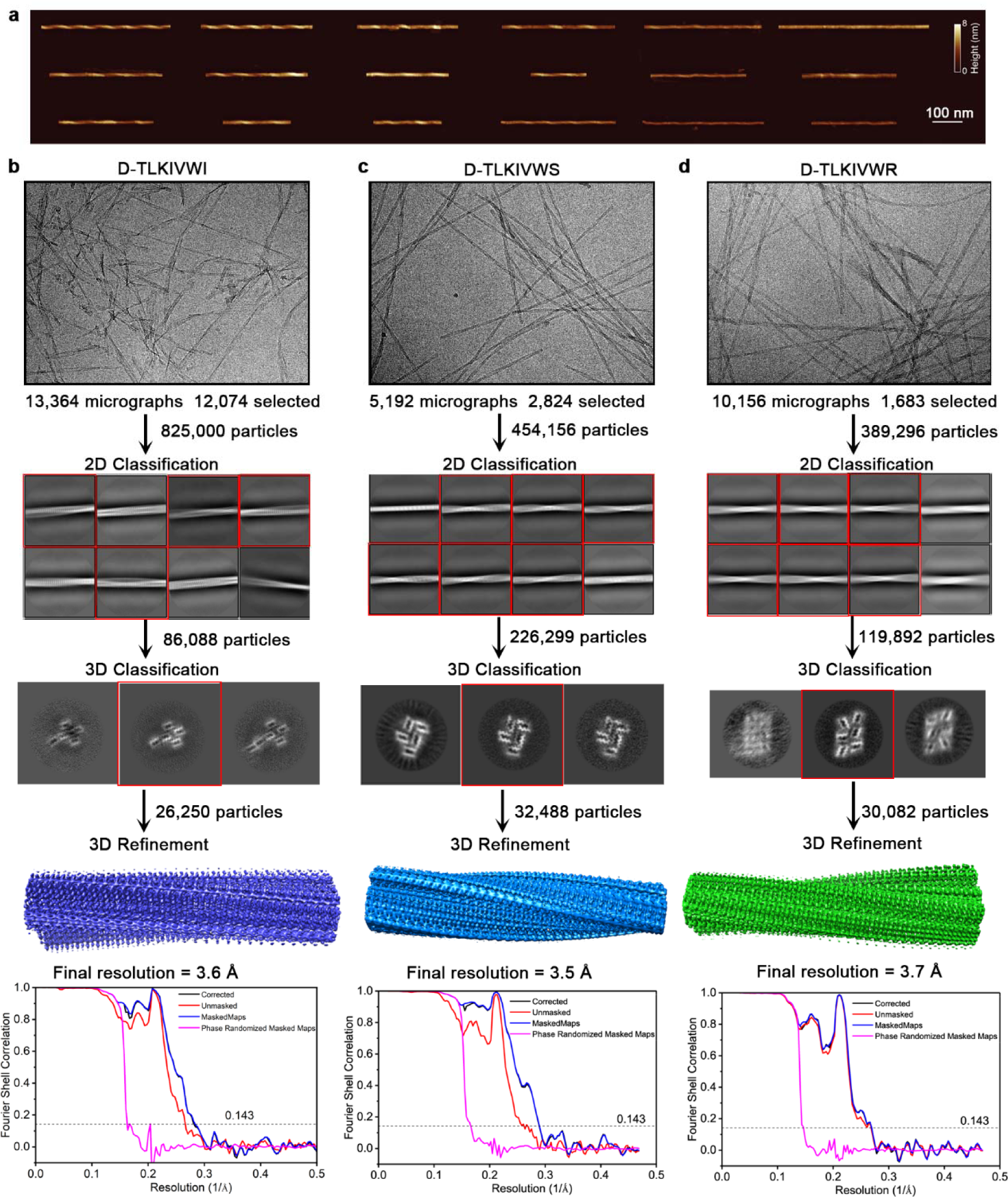
AFM and Cryo-EM studies of D-TLKIVWX (X = I, S and R) amyloid fibrils. **a**, AFM images of 18 polymorphs of D-TLKIVWI fibrils. Scale bar, 100 nm. **b-d**, Cryo-EM imaging and data analysis of (**b**) D-TLKIVWI fibrils, (**c**) D-TLKIVWS fibrils and (**d**) D-TLKIVWR fibrils. Notice that all three peptide fibrils have a right-handed twist.

**Extended Data Fig. 7.**
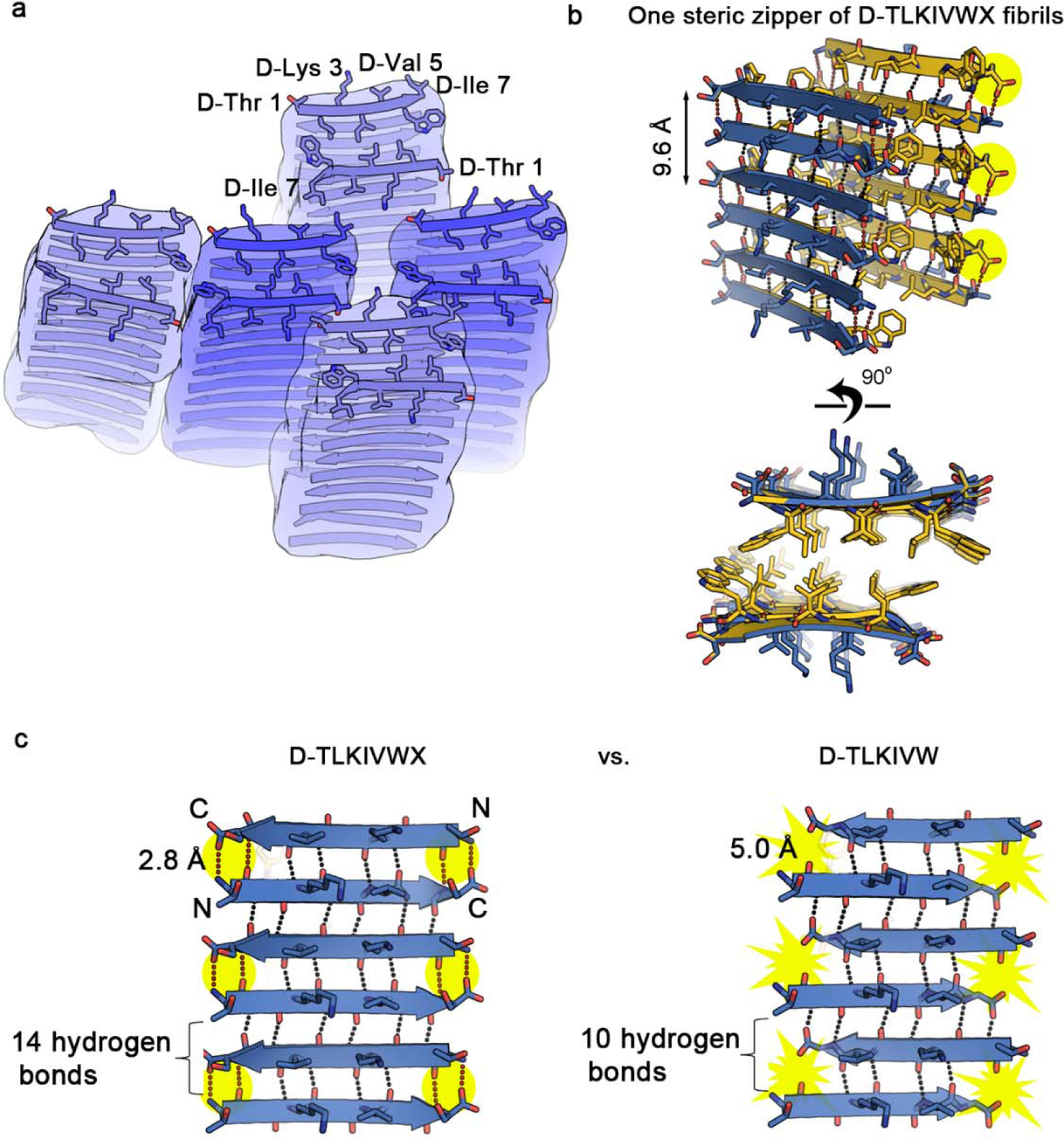
Structural explanation of why the absence of a seventh residue in D-TLKIVW prohibits its fibril formation. **a**, Schematic model of the amyloid-like fibrils formed by D-TLKIVWI**. b**, Views perpendicular to the fibril axis (upper row) and down the fibril axis (lower row) of one steric zipper of D-TLKIVWX amyloid-like fibrils. **c**, D-TLKIVW fails to form stable amyloid fibrils like D-TLKIVWX due to its reduced number of backbone hydrogen bonds (10 vs. 14) and the increased distance between the negatively charged C-terminal carboxyl group and the positively charged N-terminal amine group of the adjacent strand (5.0 Å vs. 2.8 Å), as compared between the left- and right-side panel c.

**Extended Data Fig. 8.**
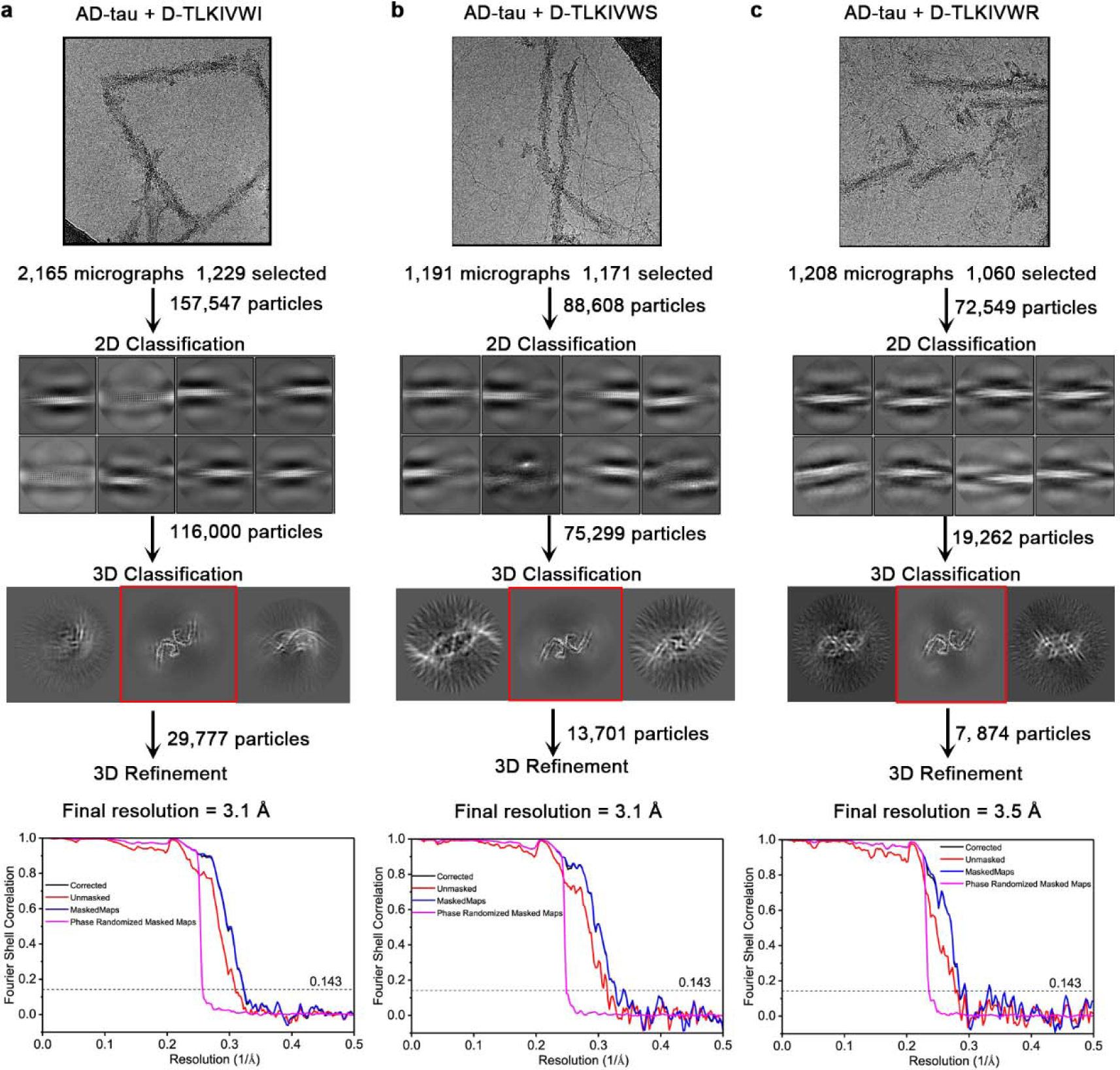
Cryo-EM data analysis workflow of the three complexes D-TLKIVWX (X = I, S and R)-disassembling AD-tau fibrils. **a-c**, Cryo-EM imaging and data analysis of (**a**) D-TLKIVWI-, (**b**) D-TLKIVWS-, and (**c**) D-TLKIVWR-disassembling AD-tau fibrils.

**Extended Data Fig. 9.**
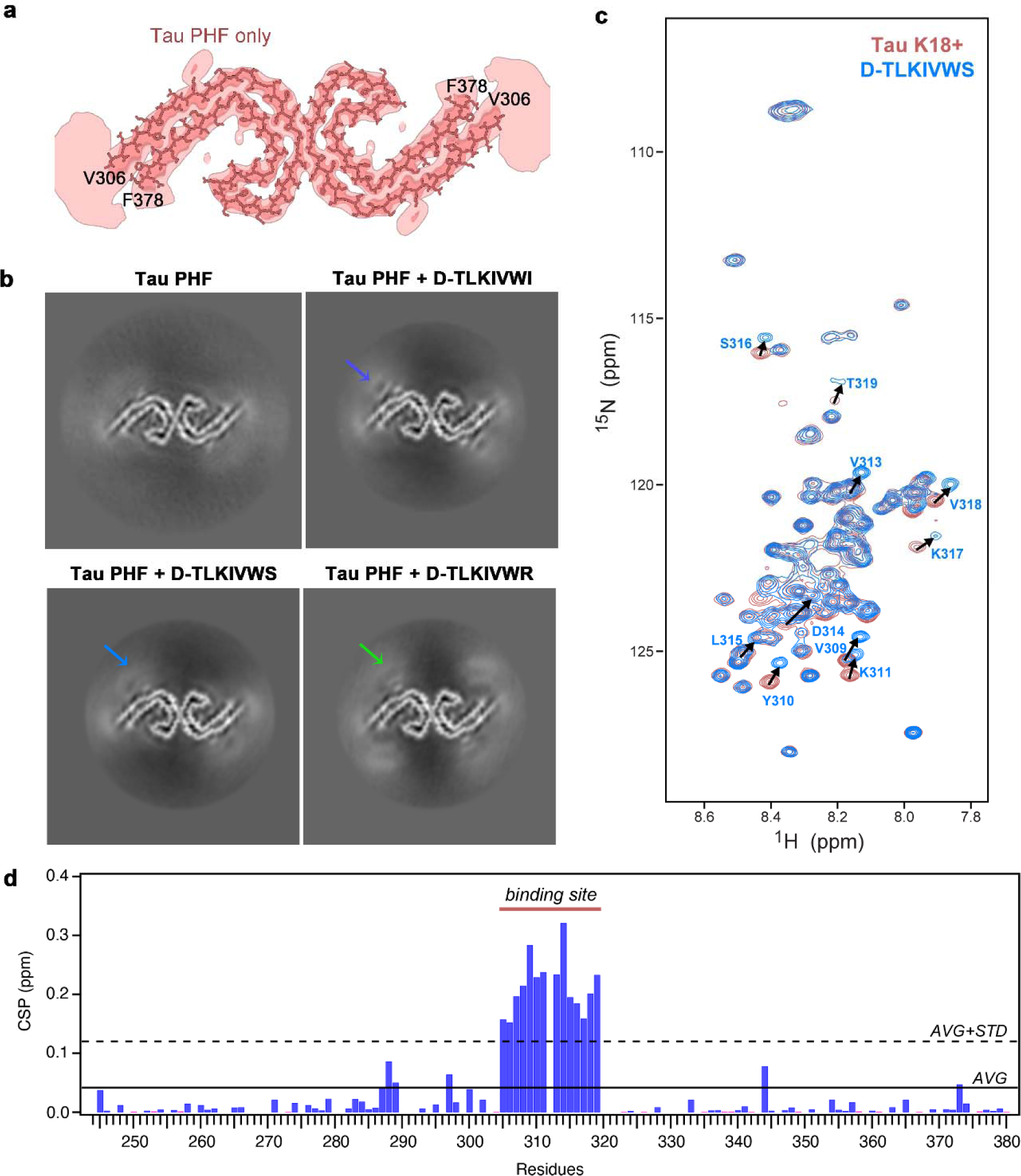
D-TLKIVWX (X = I, S and R) binds to the Val313–Thr319 residues of tau PHF. **a,** Cross-sectional cryo-EM density map and atomic models of tau PHF. A sharpened, high-resolution map is shown in dark salmon. Unsharpened, a 7 Å low pass filtered density map is shown in light salmon. **b**, 7 Å low-pass filtered cryo-EM maps of tau PHF control, and D-TLKIVWX (X = I, S, R)-disassembling tau PHF. The density map interpreted as peptide steric zippers is indicated by arrows. **c**, ^1^H-^15^N-HSQC spectral overlay of free ^15^N,^13^C-labeled tau K18+ (salmon) and with 10-fold molar excess of D-TLKIVWS peptide after incubating for 1 month at 4 L (marine). Representative residues with significant chemical shift changes are labeled and resonance shifts indicated with arrows. **d**, Chemical shift perturbation between free and D-TLKIVWI peptide bound ^15^N,^13^C-labeled tau K18+ monomer resonances.

**Extended Data Fig. 10.**
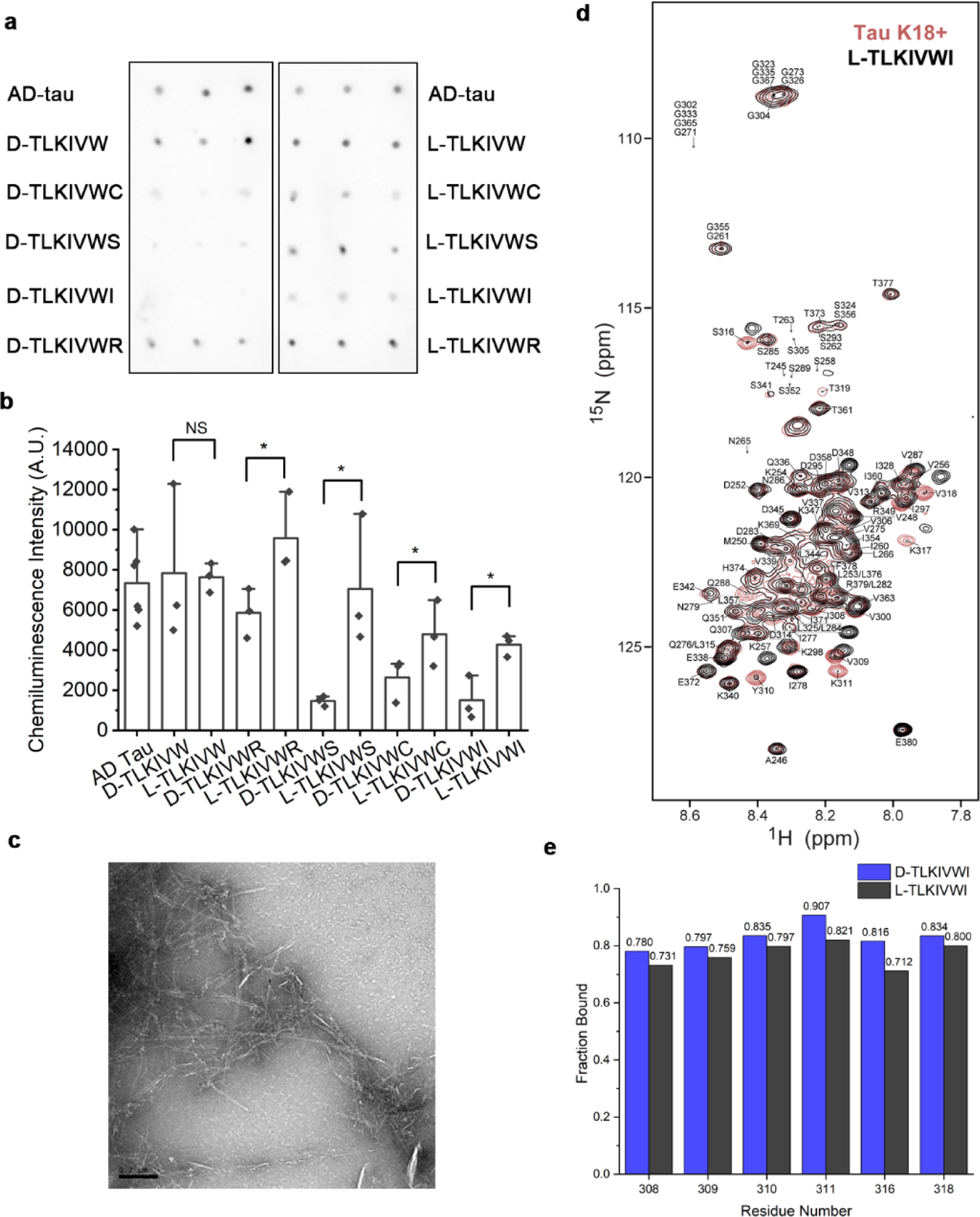
L-TLKIVWX peptides display diminished efficacy in disassembling AD-tau fibrils compared to their enantiomers D-TLKIVWX. **a**, Dot blot staining of AD-tau fibrils before and after incubation with 500 μM D/L-TLKIVW, D/L-TLKIVWX (X= C, S, I, R) at 37 L for 48 hours, probed by the GT38 antibody. Each sample was analyzed in triplicate. **b**, Quantification of the level of AD-tau fibrils in panel a by ImageJ. **c**, Negative stain TEM image of 500 μM L-TLKIVWI after incubation in 20 mM Tris-HCl, pH 7.4, 100 mM NaCl for 48 hours. Scale bar, 0.2 μm. **d**, ^1^H-^15^N-HSQC spectral overlay of free ^15^N,^13^C-labeled tau K18+ (salmon) and with 10-fold molar excess of L-TLKIVWI peptide after 2.5 months of incubation at 4 °C (black). All residues with backbone chemical shift assignments are labeled. **e**, Comparison of fraction of tau K18+ that are bound by L- and D-TLKIVWI measured by peak intensities of representative residues (I308, V309, Y310, K311, S316, and V318). Both bound and residual free tau peak intensities were from the peptide bound spectra of tau K18+ in panel d and Fig. 4d. The slightly lower efficacy of L-TLKIVWI binding tau monomers was possible due to the slow exchange regime on NMR chemical exchange timescale.

