## Supplementary material for "How short peptides can disassemble ultra-stable tau fibrils extracted from Alzheimer’s disease brain by a strain-relief mechanism": Cryo-EM statistics of D-TLKIVWX (X = I, S, R) fibrils

**Cryo-EM data collection, refinement and validation statistics**

| **Continued** | **D-TLKIVWI**  **(EMDB-44181)**  **(PDB 9B4I)** | **D-TLKIVWS**  **(EMDB-44182)**  **(PDB 9B4J)** | **D-TLKIVWR**  **(EMDB-44183)**  **(PDB 9B4K)** |
| --- | --- | --- | --- |
| **Data collection and processing** |  |  |  |
| **Scope** | Krios | Krios | Krios |
| **Energy Filter** | Selectris X | BioQuantum | BioQuantum |
| **Camera** | Falcon 4i | K3 | K3 |
| **Automation** | SerialEM | Leginon | Leginon |
| **Magnification** | x130,000 | x81,000 | x81,000 |
| **Voltage (kV)** | 300 | 300 | 300 |
| **Electron exposure (e–/Å^2^)** | 55 | 60 | 60 |
| **Defocus range (μm)** | 1.0-3.8 | 0.8-3.5 | 0.7-3.1 |
| **Pixel size (Å)** | 0.94 | 1.067 | 1.067 |
| **Symmetry imposed** | C1 | C1 | C1 |
| **Helical rise** | 9.66 | 9.56 | 9.56 |
| **Helical twist** | 1.60 | 3.50 | 2.55 |
| **Initial particle images (no.)** | 825,000 | 454,000 | 389,000 |
| **Final particle images (no.)** | 26,250 | 32,435 | 30,029 |
| **Map resolution (Å)** | 3.60 | 3.48 | 3.75 |
| **FSC threshold** | 0.143 | 0.143 | 0.143 |
| **Map resolution range (Å)** | 200-3.6 | 200-3.5 | 200-3.7 |
| **Refinement** |  |  |  |
| **Initial model used (PDB code)** | N/A | N/A | N/A |
| **Model resolution (Å)**  **FSC threshold** | 4.25  0.5 | 3.79  0.5 | 4.11  0.5 |
| **Model resolution range (Å)** | 200-3.60 | 200-3.50 | 200-3.70 |
| **Map sharpening *B* factor (Å^2^)** | 143.53 | 124.08 | 189.47 |
| **Model composition**  **Non-hydrogen atoms**  **D-peptide residues**  **Ligands** | 3720  3720  N/A | 3600  3600  N/A | 3380  3380  N/A |
| ***B* factors (Å^2^)**  **D-peptide**  **Ligand** | 17.8  N/A | 83.0  N/A | 61.4  N/A |
| **R.m.s. deviations**  **Bond lengths (Å)**  **Bond angles (°)** | 0.003  0.77 | 0.007  0.99 | 0.006  0.54 |
| **Validation**  **MolProbity score**  **Clashscore**  **Poor rotamers (%)** | 2.8  14.3  0.0 | 2.9  16.5  0.0 | 2.5  10.9  15.1 |
| **Ramachandran plot**  **Favored (%)**  **Allowed (%)**  **Disallowed (%)** | 100.0  0.0  0.0 | 100.0  0.0  0.0 | 100.0  0.0  0.0 |
