## Supplementary material for "How short peptides can disassemble ultra-stable tau fibrils extracted from Alzheimer’s disease brain by a strain-relief mechanism": Cryo-EM statistics of AD-tau and AD-tau-D-TLKIVWX (X = I, S, R) complex

**Cryo-EM data collection, refinement and validation statistics**

|  | **Tau-D-TLKIVWI**  **(EMDB-44184)**  **(PDB 9B4L)** | **Tau-D-TLKIVWS**  **(EMDB-44185)**  **(PDB 9B4M)** | **Tau-D-TLKIVWR**  **(EMDB-44186)**  **(PDB 9B4N)** | **Tau-Control**  **(EMDB-44187)**  **(PDB 9B4O)** |
| --- | --- | --- | --- | --- |
| **Data collection and processing** |  |  |  |  |
| **Scope** | Krios | Krios | Krios | Krios |
| **Energy Filter** | Selectris X | Selectris X | Selectris X | BioQuantum |
| **Camera** | Falcon 4i | Falcon 4i | Falcon 4i | K3 |
| **Automation** | SerialEM | SerialEM | SerialEM | EPU |
| **Magnification** | x130,000 | x130,000 | x130,000 | x81,000 |
| **Voltage (kV)** | 300 | 300 | 300 | 300 |
| **Electron exposure (e–/Å^2^)** | 50 | 50 | 50 | 50 |
| **Defocus range (μm)** | 0.8-4.3 | 0.9-3.9 | 0.6-2.6 | 0.4-4.0 |
| **Pixel size (Å)** | 0.94 | 0.94 | 0.94 | 1.1 |
| **Symmetry imposed** | C1 | C1 | C1 | C1 |
| **Helical rise** | 2.41 | 2.41 | 2.41 | 2.48 |
| **Helical twist** | 179.44 | 179.45 | 179.46 | 179.45 |
| **Initial particle images (no.)** | 158,000 | 89,000 | 72,000 | 105,000 |
| **Final particle images (no.)** | 26,975 | 13,648 | 7,821 | 14,048 |
| **Map resolution (Å)** | 3.03 | 3.05 | 3.53 | 3.52 |
| **FSC threshold** | 0.143 | 0.143 | 0.143 | 0.143 |
| **Map resolution range (Å)** | 200-3.0 | 200-3.0 | 200-3.5 | 200-3.5 |
| **Refinement** |  |  |  |  |
| **Initial model used (PDB code)** |  |  |  |  |
| **Model resolution (Å)**  **FSC threshold** | 3.32  0.5 | 3.26  0.5 | 3.75  0.5 | 3.81  0.5 |
| **Model resolution range (Å)** | 3.10 | 3.10 | 3.50 | 3.50 |
| **Map sharpening *B* factor (Å^2^)** | 67.32 | 47.72 | 95.15 | 120.04 |
| **Model composition**  **Non-hydrogen atoms**  **Tau**  **D-peptide** | 7044  1488 | 5870  0 | 5870  0 | 5870  0 |
| ***B* factors (Å^2^)**  **Tau**  **D-peptide** | 89.9  110.0 | 109.4  N/A | 101.3  N/A | 101.0  N/A |
| **R.m.s. deviations**  **Bond lengths (Å)**  **Bond angles (°)** | 0.008  0.91 | 0.006  0.55 | 0.006  0.55 | 0.006  0.77 |
| **Validation**  **MolProbity score**  **Clashscore**  **Poor rotamers (%)** | 2.92  15.4  7.5 | 1.79  5.0  0.0 | 2.57  9.6  4.5 | 1.85  4.7  0.0 |
| **Ramachandran plot**  **Favored (%)**  **Allowed (%)**  **Disallowed (%)** | 89.3  10.7  0.0 | 90.7  9.3  0.0 | 89.3  10.7  0.0 | 88.0  12.0  0.0 |
